# mbImpute: an accurate and robust imputation method for microbiome data

**DOI:** 10.1101/2020.03.07.982314

**Authors:** Ruochen Jiang, Wei Vivian Li, Jingyi Jessica Li

## Abstract

Microbiome studies have gained increased attention since many discoveries revealed connections between human microbiome compositions and diseases. A critical challenge in microbiome research is that excess non-biological zeros distort taxon abundances, complicate data analysis, and jeopardize the reliability of scientific discoveries. To address this issue, we propose the first imputation method, mbImpute, to identify and recover likely non-biological zeros by borrowing information jointly from similar samples, similar taxa, and optional metadata including sample covariates and taxon phylogeny. Comprehensive simulations verified that mbImpute achieved better imputation accuracy under multiple measures than five state-of-the-art imputation methods designed for non-microbiome data. In real data applications, we demonstrate that mbImpute improved the power and reproducibility of identifying disease-related taxa from microbiome data of type 2 diabetes and colorectal cancer.

## Introduction

Microbiome studies explore the collective genomes of microorganisms living in a certain environment such as soil, sea water, animal skin, and human gut. A large number of studies have confirmed the importance of microbiomes in both natural environment and human bodies [1]. For example, new discoveries have revealed the important roles microbiomes play in complex diseases such as obesity [2], diabetes [3], pulmonary disease [4, 5], and cancers [6]. These studies have shown the potential of using human microbiomes as biomarkers for disease diagnosis or therapeutic targets for disease treatment [7].

The development of high-throughput sequencing technologies has advanced microbiome studies in the last decade [8]. Microbiome studies primarily use two sequencing technologies: the 16S ribosomal RNA (rRNA) amplicon sequencing and the shotgun metagenomics sequencing. The former specifically sequences 16S rRNAs, which can be used to identify and distinguish microbes [9]. The 16S sequencing reads are either clustered into operational taxonomic units (OTUs) [10] or mapped to amplicon sequence variants (ASVs) for a higher resolution [11, 12]. The latter, often referred to as whole-genome sequencing (WGS), sequences all DNAs in a microbiome sample, including whole genomes of microbial species and host DNAs [10, 13–19], and its sequencing reads are mapped to known microbiome genome databases to quantify the abundance of each microbial species. Despite the vast differences between these two technologies, 16S and WGS data can both be processed into a similar data format about abundances of microbes in biological samples: a taxon count matrix with rows as samples (which often correspond to subjects) and columns as taxa (i.e., OTUs for 16S rRNA data and species for WGS data), and each entry corresponds to the number of reads mapped to a taxon in a sample. It is worth noting that the total read count per sample, i.e., the sum of entries in a row of the count matrix, differs by five orders of magnitude between the two technologies: ∼ 10^3^ per sample for 16S rRNA data and ∼ 10^8^ for WGS data [20].

A critical challenge in microbiome data analysis is the existence of excess zeros in taxon counts, a phenomenon prevalent in both 16S rRNA and WGS data [20]. The excess zeros belong to three categories by origin: biological, technical, and sampling zeros [21]. Biological zeros represent true zero abundances of non-existent taxa in samples. In contrast, both technical and sampling zeros are non-biological zeros with different origins: technical zeros arise from presequencing experimental artifacts (e.g., DNA degradation during library preparation and inefficient sequence amplification due to factors such as GC content bias) [22], while sampling zeros are due to limited sequencing depths. Although WGS data have much larger per-sample total read counts than 16S data have, they still suffer from excess zeros because they sequence more nucleic acid sequences (microbial genomes instead of 16S rRNAs) and widespread host DNA contamination reduces the effective sequencing depths for microbial genomes [23–25].

This data sparsity issue has challenged the statistical analysis of microbiome data, as most state-of-the-art methods have poor performance on data containing too many zeros. Adding a pseudo-count of one to zeros is a common, simple approach [26, 27], but it is known to be ad-hoc and suboptimal as it cannot not distinguish biological zeros from technical and sampling zeros [28, 29]. Kaul et al. [30] developed an approach to distinguish these three types of zeros and only correct the sampling zeros; however, their correction is still a simple addition of a pseudo-count of one, ignoring the fact that the (unobserved) actual counts of these sampling zeros may not be exactly one.

In particular, this data sparsity issue has greatly hindered the differentially abundant (DA) taxon analysis, which is to identify the taxa that exhibit significantly different abundances between two groups of samples [13]. Microbiome researchers employ two major types of statistical methods to identify DA taxa. Methods of the first type are based on parametric models [7, 26, 31–38]. For example, the zero-inflated negative Binomial generalized linear model (ZINB-GLM) is used in [7, 31, 32], the DESeq2-phyloseq method uses the negative Binomial regression [33, 34], and the metagenomeSeq method uses the zero-inflated Gaussian model [35]. However, the different parametric model assumptions do not always fit data well [39]. Methods of the second type perform non-parametric statistical tests that do not assume specific distributions, and widely-used methods include the Wilcoxon rank-sum test [14–19] and ANCOM [27]. A major drawback of these non-parametric methods is that a taxon would be called DA if its zero proportions differ significantly between two groups of samples, but this difference is unlikely biologically meaningful due to the prevalence of technical and sampling zeros. Both types of methods consider taxon abundances at one of three scales: counts [7, 31, 32, 34], log-transformed counts [35], and proportions (i.e., each taxon’s count is divided by the total of all taxa’s counts in a sample) [26, 27, 36–38]. We note that excess zeros would negatively affect taxon abundances at all the three levels.

In addition to DA taxon analysis, other microbiome data analyses, such as the construction of taxon interaction networks [40–43], are also impeded by the data sparsity challenge. If using the zero-inflated modeling approach, each task calls for a specialized model development, which is often complicated or unrealistic for most microbiome researchers. Hence, a flexible and robust approach is needed to address the data sparsity issue for microbiome research.

Imputation is a widely-used technique to recover missing data and facilitate data analysis. It has various successful applications in many fields such as recommender systems (e.g., the Netflix challenge [44]), image and speech reconstruction [45–47], imputation of unmeasured epigenomics datasets [48], missing genotype prediction in genome-wide association studies [49], and the more recent gene expression recovery in single-cell RNA-sequencing (scRNA-seq) data analysis [50–54]. Microbiome and scRNA-seq data have similar count matrix structures if one considers samples and taxa as analogs to cells and genes, and both data have excess non-biological zeros. Given the successes of scRNA-seq imputation methods, it is reasonable to hypothesize that imputation will also relieve the data sparsity issue in microbiome data. Although there are methods utilizing matrix completion in the microbiome field, their main purpose is to perform community detection or dimension reduction instead of imputation [55, 56]. Two distinct features of microbiome data make direct application of existing imputation methods suboptimal. First, microbiome data are often accompanied by metadata including sample covariates and taxon phylogeny, which, however, cannot be used by existing imputation methods. In particular, phylogenetic information is known to be valuable for microbiome data analysis [57–64], as taxa closely related in a phylogeny are likely to have similar functions and abundances in samples [65–68]. Second, microbiome data has a much smaller number of samples (often in hundreds) than the number of cells (often in tens of thousands) in scRNA-seq data, making those deep-learning based imputation methods inapplicable [54, 69]. On the other hand, the smaller sample size allows microbiome data to afford an imputation method that focuses more on imputation accuracy than computational time.

Here we propose mbImpute, the first imputation method designed for microbiome data, including both 16S and WGS data. mbImpute identifies and corrects zeros and low counts that are unlikely biological (for ease of terminology, we will refer to them as non-biological zeros in the following text) in microbiome taxon count data. The goal of mbImpute is to provide a principled data-driven approach to relieve the data sparsity issue due to excess non-biological zeros. To achieve this, mbImpute leverages three sources of information: a taxon count matrix, sample covariates (e.g., sample library size and subjects’ age, gender, and body mass index), and taxon phylogeny, with the latter two sources optional. mbImpute takes a two-step approach (Fig. 1): it first identifies likely non-biological zeros and second imputes them by borrowing information from similar taxa (determined by both phylogeny and counts), similar samples (in terms of taxon counts), and sample covariates if available (see illustration of the imputation step in Supplementary Fig. S1). The imputed data are expected to contain recovered taxon counts and thus facilitate various downstream analyses, such as the identification of DA taxa and the construction of taxon interaction networks. Microbiome researchers can use mbImpute to avoid the hassles of handling excess zeros in individual analysis tasks and to enjoy the flexibility of building up data analysis pipelines.

**Figure 1:**
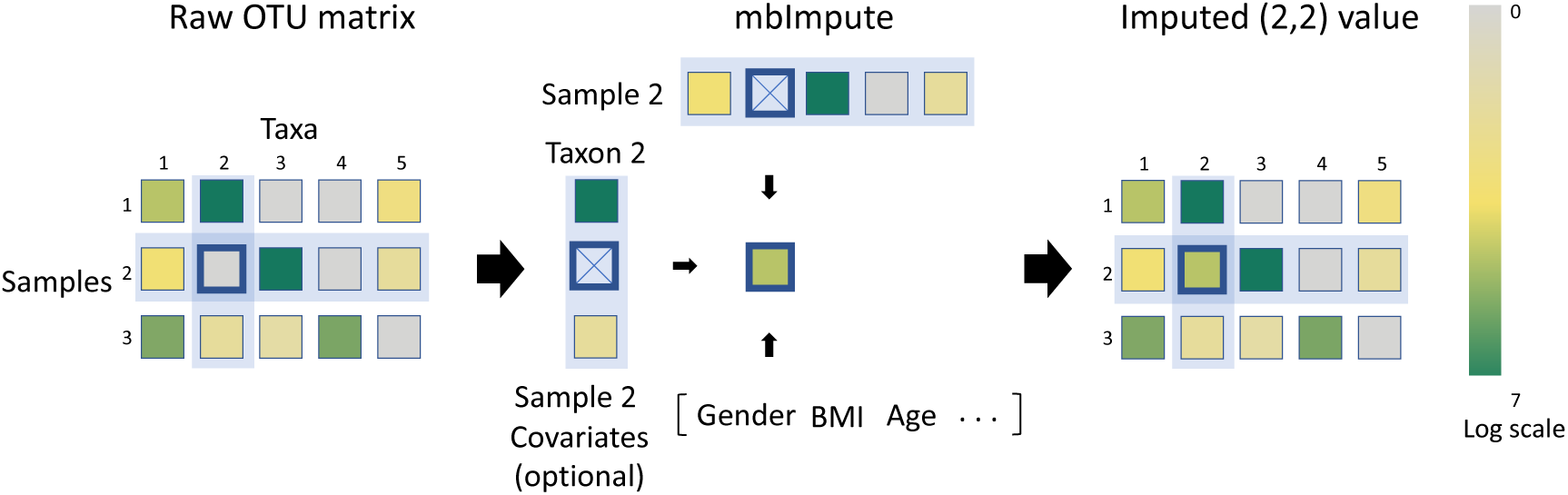
An illustration of mbImpute. After mbImpute identifies likely non-biological zeros, it imputes them (e.g. the abundance of taxon 2 in sample 2) by jointly borrowing information from similar samples, similar taxa, and sample covariates if available (details in Methods).

## Results

### mbImpute outperforms non-microbiome imputation methods in recovering missing taxon abundances and empowering DA taxon identification

As there are no imputation methods for microbiome data, we benchmarked mbImpute against five state-of-the-art imputation methods designed for non-microbiome data, including four popular scRNA-seq imputation methods (scImpute [50], SAVER [52], MAGIC [51], and ALRA [53]) and a widely-used general imputation method softImpute [70]. We designed two simulation studies, and the common goal was to obtain a “complete” microbiome dataset without non-biological zeros, so that imputation accuracy could be evaluated by comparing the imputed data with the complete data. The first study simulated complete data from a generative model that was fitted to a real WGS dataset of type 2 diabetes (T2D) samples [18], and the second, more realistic simulation study took a sub-dataset with fewer than 15% zeros as the complete data from another real WGS dataset of T2D samples [19]. In both simulation studies (see Supplementary), non-biological zeros were introduced into the complete data by mimicking the observed zero patterns in real datasets, resulting in the zero-inflated data. After applying the six methods to the zero-inflated data in both studies, we compared these methods’ imputation accuracy in three aspects: (1) the mean squared error (MSE) between the imputed data and the complete data, (2) the Pearson correlation between each taxon’s abundances in the imputed data and those in the complete data, and (3) the Wasserstein distance between the distribution of taxa’s abundance means and standard deviations in the imputed data and that in the complete data. Fig. 2a–d illustrate the comparison results, which indicate that mbImpute achieves the best overall performance in all the three aspects. In particular, Fig. 2c–d and Supplementary Fig. S2 show that the imputed data by mbImpute best resemble the complete data, verifying the advantage of mbImpute in recovering missing taxon abundances in microbiome data.

**Figure 2:**
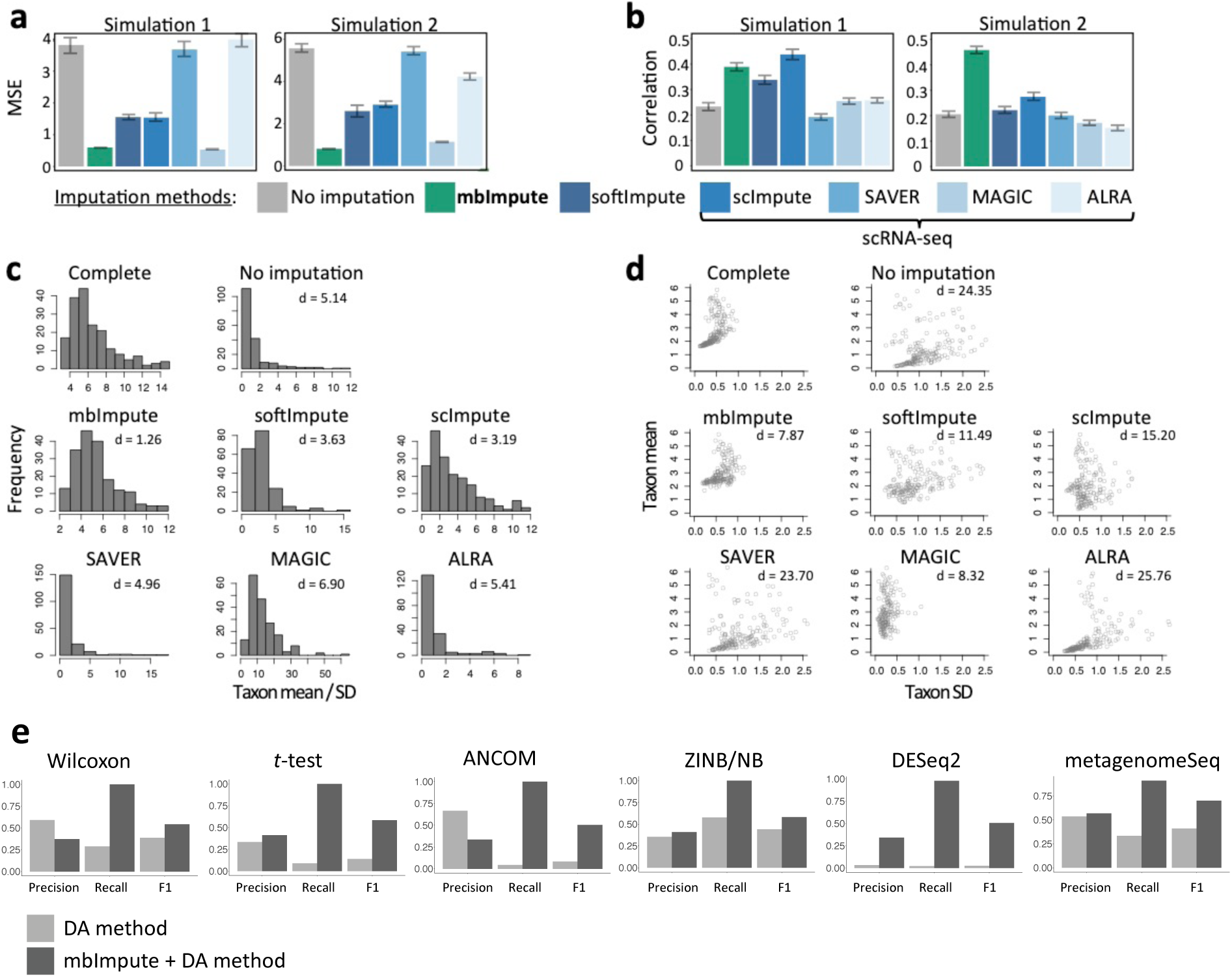
mbImpute outperforms state-of-the-art imputation methods designed for non-microbiome data and enhances the identification of DA taxa. **(a)** Mean squared error (MSE) and **(b)** mean Pearson correlation of taxon abundances between the complete data and the zero-inflated data (“No imputation,” the baseline) or the imputed data by each imputation method (mbImpute, softImpute, scImpute, SAVER, MAGIC, and ALRA) in Simulations 1 and 2 (see Supplementary). **(c)-(d)** For each taxon, the mean and standard deviation (SD) of its abundances were calculated for the complete data, the zero-inflated data, and the imputed data by each imputation method in Simulation 1; (c) shows the distributions of the taxon mean / SD and the Wasserstein distance between every distribution and the complete distribution; (d) shows the taxa in two coordinates, mean vs. SD, and the Euclidean distance between the taxa in every (zero-inflated or imputed) dataset and the complete data in these two coordinates. **(e)** Accuracy (Precision, recall and F1 scores) of six DA methods (Wilcoxon rank-sum test, *t*-test, ANCOM, ZINB/NB-GLM, DESeq2-phyloseq, metagenomeSeq) on raw data (light color) and imputed data by mbImpute (dark color) in Simulation 4.

We next demonstrated that mbImpute is a robust method. The core of mbImpute is to borrow three-way information from similar samples, similar taxa, and sample covariates to impute non-biological zeros in microbiome data (see Methods). In the aforementioned second simulation study, we broke up similar samples in the real T2D WGS data when we selected the complete data, a situation not optimal for mbImpute; however, mbImpute still outperforms existing imputation methods (Fig. 2a–b). To further test for the robustness of mbImpute, we designed a third simulation study including four simulation schemes, where information useful for imputation was encoded in sample covariates only, samples only, taxa only, or three sources together (see Supplementary). Supplementary Fig. S3a shows that, after applied to the zero-inflated data, mbImpute effectively recovers non-biological zeros and reduces the MSE under every scheme. These results verify the robustness of mbImpute in selectively leveraging information useful for imputation.

We designed the fourth simulation to mimic a typical microbiome WGS study that aims to identify DA taxa between two sample groups. We simulated data for 300 taxa in 120 samples, 60 per group (see Supplementary). Supplementary Fig. S3b shows the two-dimensional visualization of the complete data (without non-biological zeros), zero-inflated data (with non-biological zeros), and imputed data (after mbImpute was applied to zero-inflated data). Compared with the zero-inflated data, the 120 samples are more clearly separated into two groups after imputation. We next performed the DA taxon analysis to verify that imputation can boost the power of detecting DA taxa from the zero-inflated data. Specifically, we applied three state-of-the-art DA methods: the Wilcoxon rank-sum test, ANCOM, and ZINB-GLM. Among the available DA methods, the Wilcoxon rank-sum test is the most widely-used in microbiome studies [14–19], ANCOM is one of the most-cited microbiome-specific DA method [27], and ZINB-GLM was found as the most desirable count-model-based method in a comparative study [31]. We also implemented the imputation-empowerd DA analysis: applying an imputation method to the zero-inflated data, and then identifying DA taxa from the imputed data. We included two imputation methods: mbImpute and softImpute. We chose softImpute as the benchmark imputation method in this DA analysis for two reasons: first, softImptue is a general imputation method unspecific to a particular data type; second, softImpute was observed to have good performance in the first two simulations (Fig. 2a–d). After imputation, we employed the two-sample *t*-test for DA taxon identification, because each taxon’s logarithmic transformed counts (in the complete data) follow a Normal distribution in each sample group (see Supplementary); thus, if imputation is effective, the Normal distributions should be recovered and the *t*-test should be more powerful than the Wilcoxon test. To evaluate the accuracy of DA taxon identification, we used the DA taxa detected by the *t*-test on the complete data as the ground truth. Then we calculated the precision, recall and F_1_ score of each method by comparing its detected DA taxa to the ground truth. Under the p-value threshold of 0.1 (Supplementary Fig. S3c left), the two imputation-empowered DA methods achieve better recall and F_1_ scores than the three existing DA methods. Although ANCOM has the highest precision, it has the lowest and close-to-zero recall, suggesting that it finds too few DA taxa. Between mbImpute and softImpute, results under this p-value threshold do not draw a clear conclusion: mbImpute has a better precision but a worse recall, and the two methods have similar F_1_ scores. To thoroughly compare the five methods, we plotted their performance at varying thresholds in receiver operating characteristic (ROC) curves (Supplementary Fig. S3c right), which show that mbImpute has the largest area under the curve (AUC) and outperforms the three DA methods and softImpute.

To further evaluate the performance of mbImpute on 16S rRNA sequencing data, we used a 16S simulator sparseDOSSA [71] to generate abundances of 150 taxa in 100 samples under two conditions (see Supplementary). Among these 150 taxa, 45 are predefined as truly DA taxa. We applied six existing DA methods, including the Wilcoxon rank-sum test, the two-sample *t*-test, ANCOM, ZINB/NB-GLM, DESeq2-phyloseq, and metagenomeSeq. (Note that ZINB-GLM is applied to the zero-inflated data, while NB-GLM is applied to the imputed data because the imputed data are no longer zero inflated.) To evaluate the accuracy of DA taxon identification, we calculated the precision, recall and F_1_ score of each method, with or without using mbImpute as a preceding step, by comparing each method’s detected DA taxa to the truly DA taxa. Under the p-value threshold of 0.1, the mbImpute-empowered DA methods consistently have better F_1_ scores than those of the same DA methods without imputation. In particular, mbImpute improves both precision and recall rates of four DA methods: the *t*-test, ZINB/NB-GLM, DESeq2-phyloseq, and metagenomeSeq (Fig. 2e).

### mbImpute improves the reproducibility and reliability of identifying T2D microbiome markers

To demonstrate that mbImpute can benefit the identification of DA taxa in real microbiome data, we applied the six DA methods to two T2D WGS datasets: Qin et al. and Karlsson et al., with or without using mbImpute as a preceding step. We observed that taxon abundance distributions are approximately Normal after imputation (Supplementary Fig. S4). We analyzed the identified T2D-enriched taxa in two aspects. First, we examined the overlap of these identified taxa by each method, with or without mbImpute, between the two datasets. Fig. 3a shows that mbImpute improves the reproducibility of all these DA methods, whose identified T2D-enriched taxa have increased overlaps after mbImpute is used (see Venn diagrams in Supplementary Fig. S5). Second, we investigated whether the T2D-enriched taxa identified in one dataset are reliable biomarkers for predicting T2D in another dataset. Towards this goal, we trained a random forest classifier [72] on one dataset with features as the T2D-enriched taxa identified from the other dataset. Then we calculated the 5-fold cross-validation accuracy, which reflects the reliability of the identified T2D-enriched taxa as biomarkers. Fig. 3b shows that mbImpute improves this reliability for all the DA methods but ANCOM, whose accuracy stays unchanged after mbImpute. The improvement is especially significant for the Wilcoxon rank-sum test, ZINB/NB-GLM, DESeq2-phyloseq, and metagenomeSeq. For example, the classification accuracy of the T2D-enriched taxa identified by DESeq2-phyloseq increases from 62% without mbImpute to 75% with mbImpute. As a positive control, we also evaluated the classification accuracy when no DA methods are used but random forest automatically selects predictive features from all taxa. Encouragingly, we found that the accuracy becomes comparable to the positive control when ZINB/NB-GLM and DESeq2-phyloseq are used after mbImpute. Our results demonstrate that mbImpute improves the reproducibility of DA taxon identification between two T2D datasets, and that the identified DA taxa after mbImpute have better cross-dataset predictive power.

**Figure 3:**
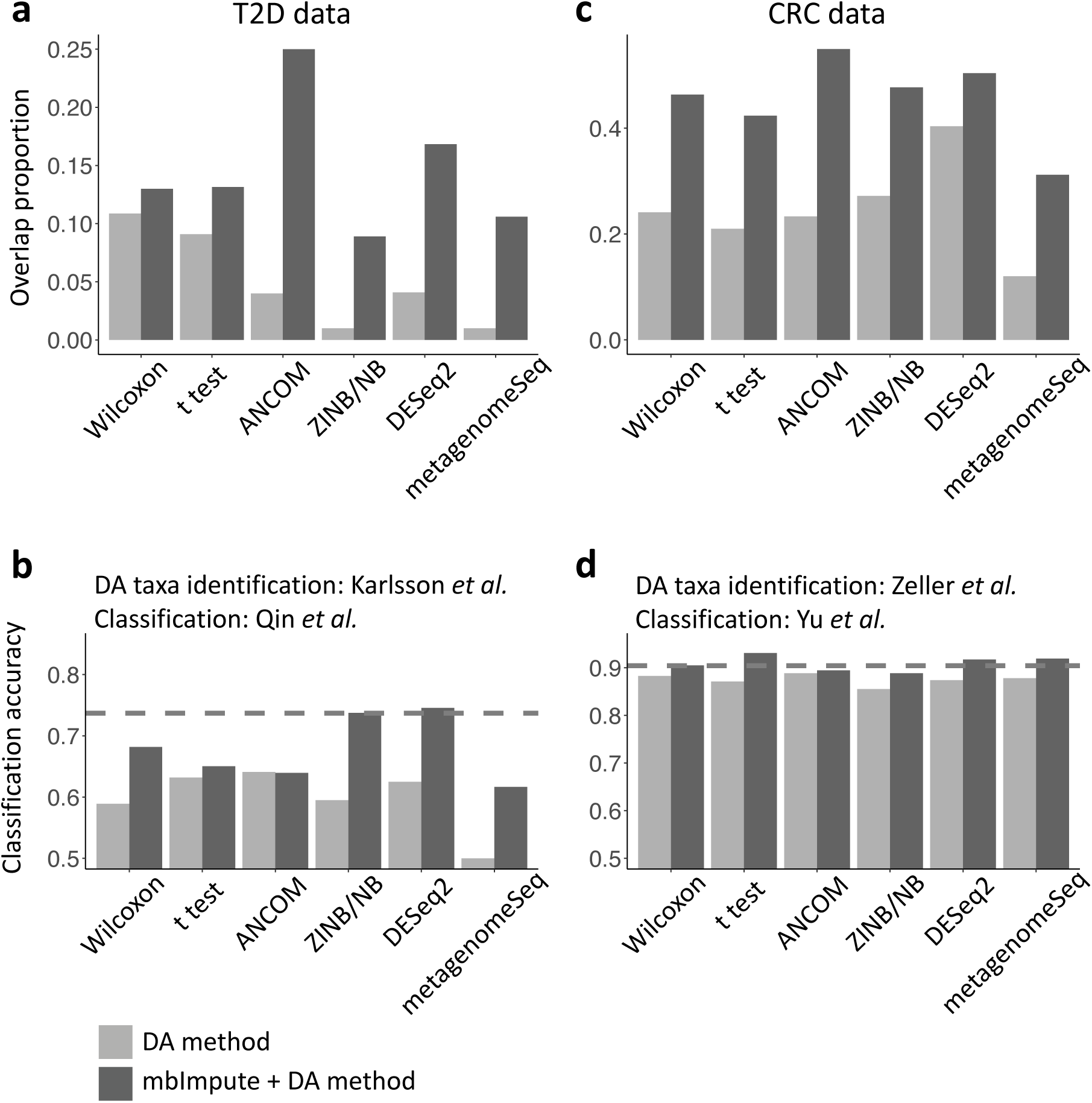
mbImpute increases the reproducibility of DA taxon identification and the accuracy of sample classification in cross-data studies. **(a)** The overlapping proportion (taxa identified as DA in both of the datasets / total number of taxa identified in either of the datasets) of identified T2D-enriched taxa between two T2D datasets [18, 19] for six DA methods, Wilcoxon rank-sum test (Wilcoxon), *t*-test, ANCOM, ZINB/NB-GLM (ZINB/NB), DESeq2-phyloseq (DESeq2), metagenomeSeq, before (light color) and after imputation (dark color). **(b)** The proportion of CRC-enriched taxa identified in at least two datasets among four CRC data [14–17] by the six DA methods before (light color) and after imputation (dark color). **(c)** The barplots show classification accuracy of prediction using random forest algorithm on the T2D status of Qin et al. by using the identified DA taxa in Karlsson et al. using six DA methods before (light color) and after imputation (dark color). The dotted horizontal line shows the prediction accuracy using random forest that automatically selects predictive features from all the taxa in Qin et al. to predict T2D statues. **(d)** The barplots show classification accuracy of prediction using random forest on the T2D status of Yu et al. by using the identified DA taxa in Zeller et al. using six DA methods before (light color) and after imputation (dark color). The dotted horizontal line shows the prediction accuracy using random forest that automatically selects predictive features from all the taxa in Yu et al. to predict CRC statues.

Further, We focused on four genera: *Streptococcus*, *Lactobacillus*, *Clostridium*, and *Actinomyces*, which have all been previously reported as enriched in T2D [73–79] (see Supplementary for the literature evidence). In these four genera, the mbImpute-empowered *t*-test discovers species-level taxa that are DA and highly enriched in T2D samples but missed by the Wilcoxon test applied to the raw data, as shown in Fig. 4a. Moreover, we observed an interesting phenomenon: some Clostridium species taxa (Fig. 4a left panel, the third genus from the top) are no longer detected as enriched in T2D samples after imputation, seemingly violating our claim that mbImpute can empower DA taxon identification as we observed in the fourth simulation. However, by examining the abundance distributions of such taxa, *Clostridium symbiosum* and *Clostridium citroniae* for example (Fig. 4a right panel top row), we found that their non-zero abundance distributions are hardly distinguishable between the T2D and control samples, suggesting that they are not informative markers for T2D. Nonetheless, the Wilcoxon test identifies them as DA in the raw data because they have different zero proportions between the T2D and control samples. This result shows that mbImpute can help reduce likely false positive DA taxa identified due to excess non-biological zeros. See Supplementary for a discussion on statistical definitions of DA taxa.

**Figure 4:**
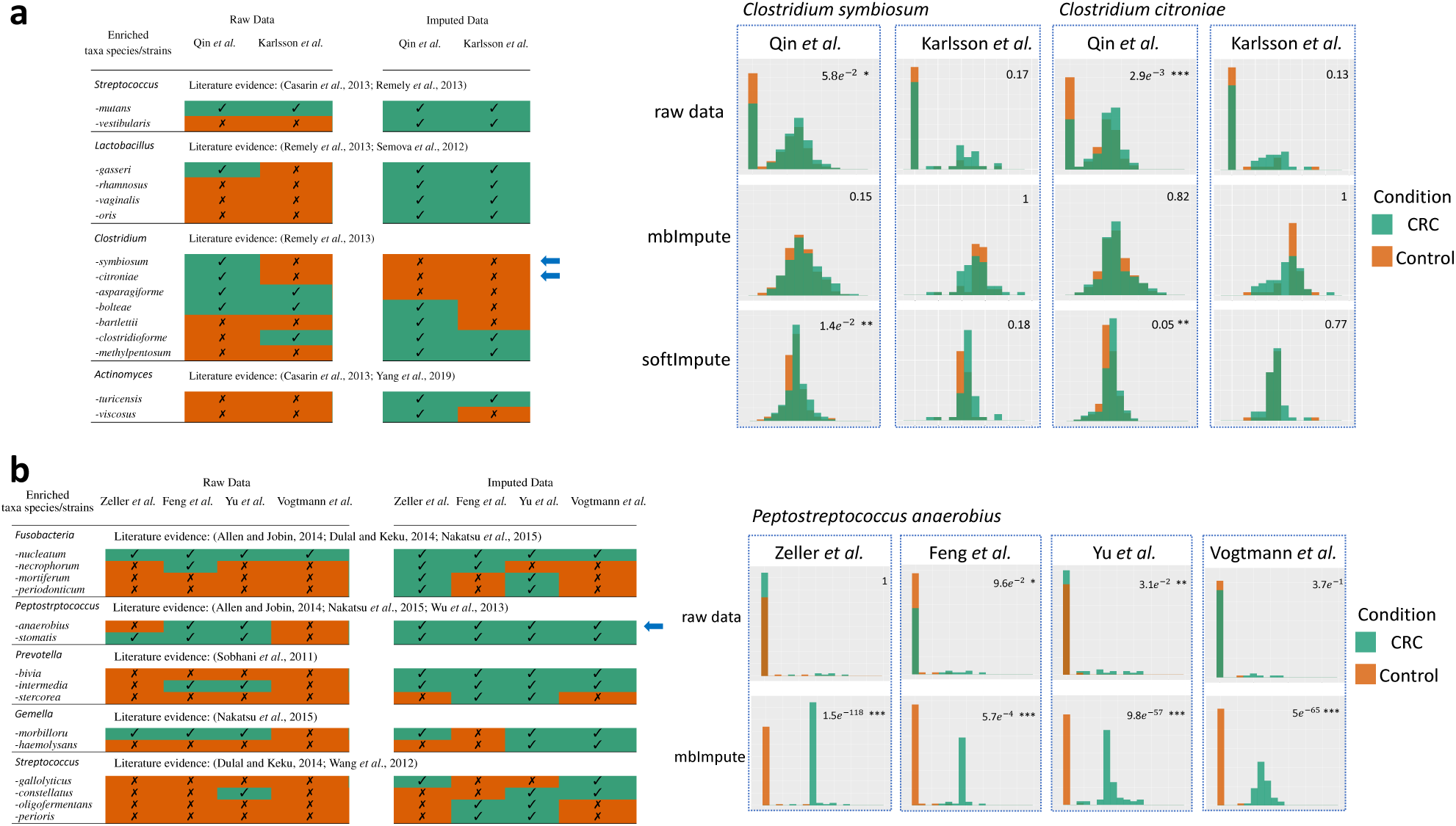
mbImpute increases the power and reproducibility of DA taxon identification in T2D and CRC WGS datasets. **(a)** Example T2D-enriched taxa identified by the Wilcoxon test on the raw data vs. the *t*-test on the imputed data by mbImpute. Literature evidence supporting the enrichment of these genera is listed. Check marks and crosses indicate the enriched and non-enriched taxa identified in each of the two datasets. Two speciess in the Clostridium genus are marked by left arrows, and their abundance (log-transformed, see Methods) distributions are plotted for the raw data (top), the imputed data by mbImpute (middle), and the imputed data by softImpute (bottom). **(b)** Example CRC-enriched taxa identified by the Wilcoxon test on the raw data vs. the *t*-test on the imputed data by mbImpute. Literature evidence supporting the enrichment of these genera is listed. Check marks and crosses indicate the enriched and non-enriched taxa identified in each of the four datasets. *Peptostreptococcus anaerobius* is marked by a left arrow, and its log-transformed abundance distributions are plotted for the raw data (top) and the imputed data by mbImpute (bottom).

We then compared mbImpute with softImpute using *Clostridium symbiosum* and *Clostridium citroniae* as examples. We observe that mbImpute retains well the distributions of non-zero abundances (Fig. 4a right panel middle row), while softImpute alters the distributions by introducing artificial spikes and shrinking the variance (Fig. 4a right panel bottom row). Such distortion of abundance distributions may mislead the DA analysis. Indeed, we found that the softImpute-empowered *t*-test identifies *Clostridium symbiosum* and *Clostridium citroniae* as DA due to the artificial distortion by softImpute. A possible reason is that softImpute is a low-rank matrix factorization method, which imputes missing matrix entries by assuming a global low-rank matrix structure. In contrast, mbImpute focuses more on local structures, i.e., how a matrix entry depends on other entries in the same row or column. The fact that mbImpute better preserves non-zero abundance distributions makes it a more reliable imputation method than softImpute for microbiome data.

### mbImpute preserves distributional characteristics of taxa’s non-zero abundances and recovers downsampling zeros

In the T2D WGS data analysis, we have found that mbImpute can well maintain the distributions of taxa’s non-zero abundances. To further verify the property of mbImpute in preserving characteristics of non-zero abundances, we examined pairwise taxon-taxon relationships in the two T2D WGS datasets: Qin et al. and Karlsson et al. For a pair of taxa, we estimated two Pearson correlations based on the raw data: one using all the samples (“raw all-sample correlation”) and the other only using the samples where both taxa have non-zero abundances (“raw non-zero-sample correlation”). We also estimated a Pearson correlation based on the imputed data by mbImpute, using all the samples (“imputed all-sample correlation”). As shown in Fig. 5, there are vast differences between the raw all-sample correlations and the corresponding raw non-zero-sample correlations. However, the imputed all-sample correlations much resemble the corresponding raw non-zero-sample correlations, suggesting that mbImpute well preserves pairwise taxon-taxon correlations encoded in taxa’s non-zero abundances.

**Figure 5:**
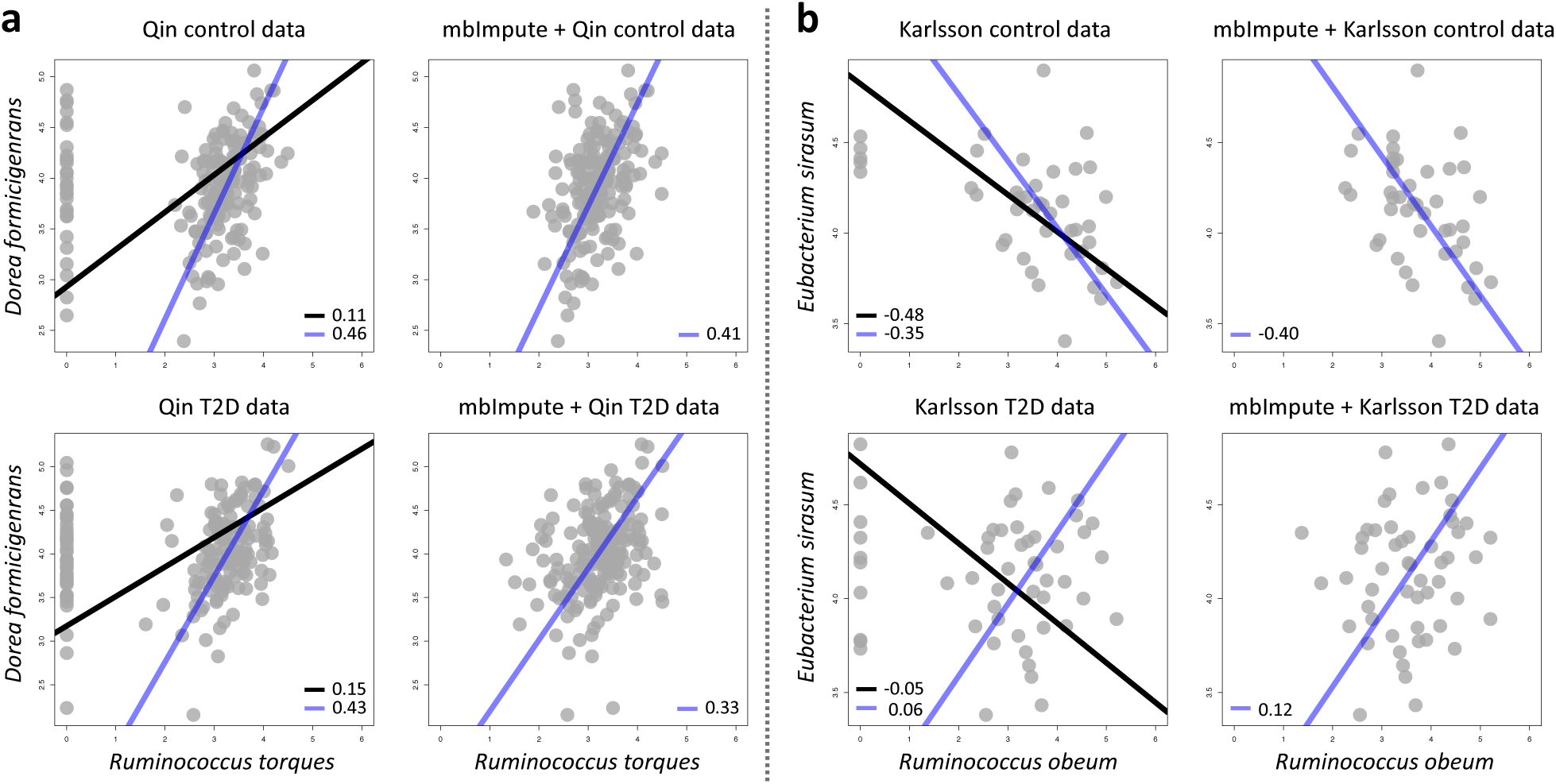
mbImpute preserves distributional characteristics of taxa’s non-zero abundances. **(a)** Top: two scatter plots show the relationship between the abundances of *Dorea formicigenerans* and *Ruminococcus torques* in Qin’s control samples, with or without using mbImpute as a preceding step. The left plot shows two standard major axis (SMA) regression lines and two corresponding Pearson correlations based on the raw data (balck: based on all the samples; blue: based on only the samples where both taxa have non-zero abundances). The right plot shows the SMA regression line (blue) and the Pearson correlation using all the samples in the imputed data. Bottom: two scatter plots for the same two taxa in ‘s T2D samples, with lines and legends defined the same as in the Top panel. **(b)** Four scatter plots show the SMA regression lines and correlations between *Eubacterium sirasum* and *Ruminococcus obeum* in ‘s control and T2D samples, with lines and legends defined the same as in (a).

We also explored the linear relationship of each taxon pair using the standard major axis (SMA) regression, which, unlike the least-squares regression, treats two taxa symmetric and considers randomness in both taxa’s abundances. For a pair of taxa, we performed two SMA regressions on the raw data: one using all the samples (“raw all-sample regression”) and the other using only the samples where both taxa have non-zero abundances (“raw non-zero-sample regression”). We also performed the SMA regression on the imputed data by mbImpute, using all the samples (“imputed all-sample regression”). Fig. 5 shows that the raw all-sample regressions and the raw non-zero-sample regressions return strongly different lines. Especially, the two lines between the two taxa *Eubacterium sirasum* and *Ruminococcus obeum* in the Karlsson et al. data (Fig. 5b bottom left) exhibit slopes of opposite signs. In contrast, the imputed all-sample regressions output lines with similar slopes to those of the raw non-zero-sample regressions. This result again confirms mbImpute’s capacity to preserve characteristics of taxa’s non-zero abundances in microbiome data.

Our results echo existing concerns about spurious taxon-taxon correlations estimated from microbiome data due to excess non-biological zeros [80, 81]. In other words, taxon-taxon correlations cannot be accurately estimated from raw data. Without imputation, an intuitive approach is to use taxa’s non-zero abundances to estimate taxon-taxon correlations; however, this approach reduces the sample size for estimating each taxon pair’s correlation, as the samples with zero abundances for either taxon would not be used, and it also makes different taxon pairs’ correlation estimates rely on different samples. To address these issues, mbImpute provides another approach: its imputed data allow taxon-taxon correlations to be estimated from all the available samples. We have verified this mbImpute approach by the fact that the correlation estimates from the imputed data resemble those from the non-zero abundances in the raw data.

In addition, based on the T2D WGS dataset generated by Qin et al., we verified mbImpute’s capacity to identify non-biological zeros generated by downsampling. In each sample (i.e., each row in the sample-by-taxon count matrix), we assigned every taxon a sampling probability proportional to its count, i.e., the larger the count, the more likely the taxon is to be sampled; based on these probabilities, we sampled 60% or 30% of the non-zero taxon counts, and we set the unsampled counts to zeros (corresponding to a removal rate of 40% or 70%). After mbImpute is applied to the downsampled count matrices, we found that mbImpute correctly identifies 95.83% and 92.83% of the newly introduced non-biological zeros under the two downsampling schemes. Before imputation, the Pearson correlations between the two downsampled matrices and the original matrix (on the log scale) are 0.76 and 0.53. After applying mbImpute to all the three matrices, the correlations are increased to 0.87 and 0.76 (Table 1). This result confirms the effectiveness of mbImpute in recovering zeros due to downsampling.

**Table 1:** Effectiveness of mbImpute in indentifying zeros due to downsampling of Qin et al.’s T2D WGS dataset. Two downsampled datasets with removal rates 40% and 70% were constructed. The first row lists the percentages of downsampling zeros identified by mbImpute; the second row lists the Pearson correlations between each of the two downsampled matrices and the original matrix (on the log scale) before imputation; the third row lists the Pearson correlations (on the log scale) after mbImpute was used. For each number, we included its error margin as the 1.96 times of the corresponding standard error over 10 replications of downsampling.

| Removal rate | 40% | 70% |
| --- | --- | --- |
| % of downsampling zeros indentified | 95.83% ± 0.46% | 92.83% ± 0.92% |
| Pearson correlation before imputation | 0.7565 ± 0.0023 | 0.5261 ± 0.0016 |
| Pearson correlation after imputation | 0.8747 ± 0.0100 | 0.7582 ± 0.0235 |

### mbImpute increases the power and reproducibility of identifying microbiome markers for colorectal cancer

Colorectal cancer (CRC) is one of the most frequently diagnosed cancer and a leading cause of cancer mortality worldwide [14, 15]. We applied the six DA methods to four CRC datasets: Zeller et al., Feng et al., Vogtmann et al., and Yu et al., with or without using mbImpute as a preceding step. We also evaluated two aspects of DA taxon identification as we did in the afore-mentioned T2D analysis: the number of identified DA taxa identified in at least two datasets and the across-dataset classification accuracy when the identified DA taxa are used as features. Fig. 3c shows that mbImpute improves the reproducibility of all these DA methods, whose identified CRC-enriched taxa have increased overlaps after mbImpute is used (see diagrams in Supplementary Fig. S6). Then we investigated whether the CRC-enriched taxa identified in one dataset are reliable biomarkers for predicting CRC in another dataset. We used the same procedures as in the T2D analysis to obtain the classification results based on random forest. Fig. 3d shows that mbImpute improves the across-dataset classification accuracy for all the six DA methods. We again set a positive control by allowing random forest to automatically select predictive features from all taxa, and we found that the accuracy of all DA methods becomes comparable to or even surpasses the positive control after mbImpute is used.

We then focused on five genera: *Fusobacterium*, *Peptostreptococcus*, *Prevotella*, *Gemella*, and *Streptococcus*, which have been previously reported as enriched in CRC [82–87] (see Supplementary for the literature evidence). In these five genera, the mbImpute-empowered *t*-test discovers species-level taxa that are DA and highly enriched in CRC samples but missed by the Wilcoxon test applied to the raw data, as shown in Fig. 4b. We use *Peptostreptococcus anaerobius* as an example to demonstrate the effectiveness of mbImpute in empowering DA taxon identification. This species taxon is identified as enriched in CRC samples by the Wilcoxon test in only two out of the four raw datasets generated by different labs. By closely examining this taxon’s abundance distributions (Fig. 4b right panel top row), we observed that non-zero abundances consistently have higher densities in the CRC samples than in the control, suggesting that this taxon should have been identified as DA in all the four datasets. The Wilcoxon test fails to identify it in ‘s and ‘s data because the dominance of zeros obscures the differences between the non-zero abundances in the CRC and control samples. However, these non-zero abundances are informative for distinguishing the CRC samples from the control; that is, if we detect this taxon with a high abundance in a patient, we should be aware of the potential implication of CRC and perform further diagnosis. mbImpute helps amplify the non-zero signals by reducing likely non-biological zeros (Fig. 4b right panel bottom row), thus empowering the identification of this taxon as DA in all the four datasets (i.e., smaller *p*-values after imputation, Fig. 4b right panel).

### mbImpute increases the similarity of microbial community structure between 16S rRNA and WGS data

We further show that mbImpute can enhance the similarity of taxon-taxon correlations inferred from micrbiome data measured by two technologies—16S rRNA sequencing and WGS. We used two microbiome datasets of healthy human stool samples: a 16S rRNA dataset from the Human Microbiome Project [88] and a WGS dataset from the control samples in Qin et al. We compared genus-level taxon-taxon correlations between these two datasets, and we did the comparison again after applying mbImpute. Fig. 6 shows that mbImpute increases the similarity between the taxon correlation structures in the two datasets. Before imputation, the Pearson correlation between the two correlation matrices (one computed from 16S rRNA taxon abundances and the other from WGS taxon abundances) is 0.59; mbImpute increases the correlation to 0.64. In particular, we observe three taxon groups (highlighted by magenta, green, and purple squares in Fig. 6) supported by both 16S rRNA and WGS data after imputation. Notably, in the magenta squares, *Acidaminococcus* has correlations with *Dialister* and *Blautia* only after imputation, and this result is consistent with the literature: *Acidaminococcus* and *Dialister* are both reported to have low abundances in healthy human stool samples [89]; *Acidaminococcus* and *Blautia* are both associated with risks of T2D and obesity, lipid profiles, and homeostatic model assessment of insulin resistance [90]. The green squares contain three bile-tolerant genera: *Alistipes*, *Bilophila*, and *Bacteroides* [91]. The raw 16S and WGS data only reveal the correlation between *Bacteroides* and *Alistipes*, but mbImpute recovers the correlations *Bilophila* has with *Alistipes* and *Bacteroides*. The purple squares indicate a strong correlation between *Sutterella* and *Prevotella* after imputation, yet this correlation is not observed in raw WGS data. We verified this correlation in the MACADAM database [92], which contains metabolic pathways associated with microbes. Out of 1260 pathways, *Sutterella* and *Prevotella* are associated with 154 and 278 pathways, respectively, and 122 pathways are in common; Fisher’s exact test finds that the overlap is statistically significant (p-value < 2.2 × 10−^16^), suggesting that *Sutterella* and *Prevotella* are indeed functionally related. Overall, our results indicate that mbImpute can facilitate meta-analysis of 16S and WGS data by alleviating the hurdle of excess non-biological zeros.

**Figure 6:**
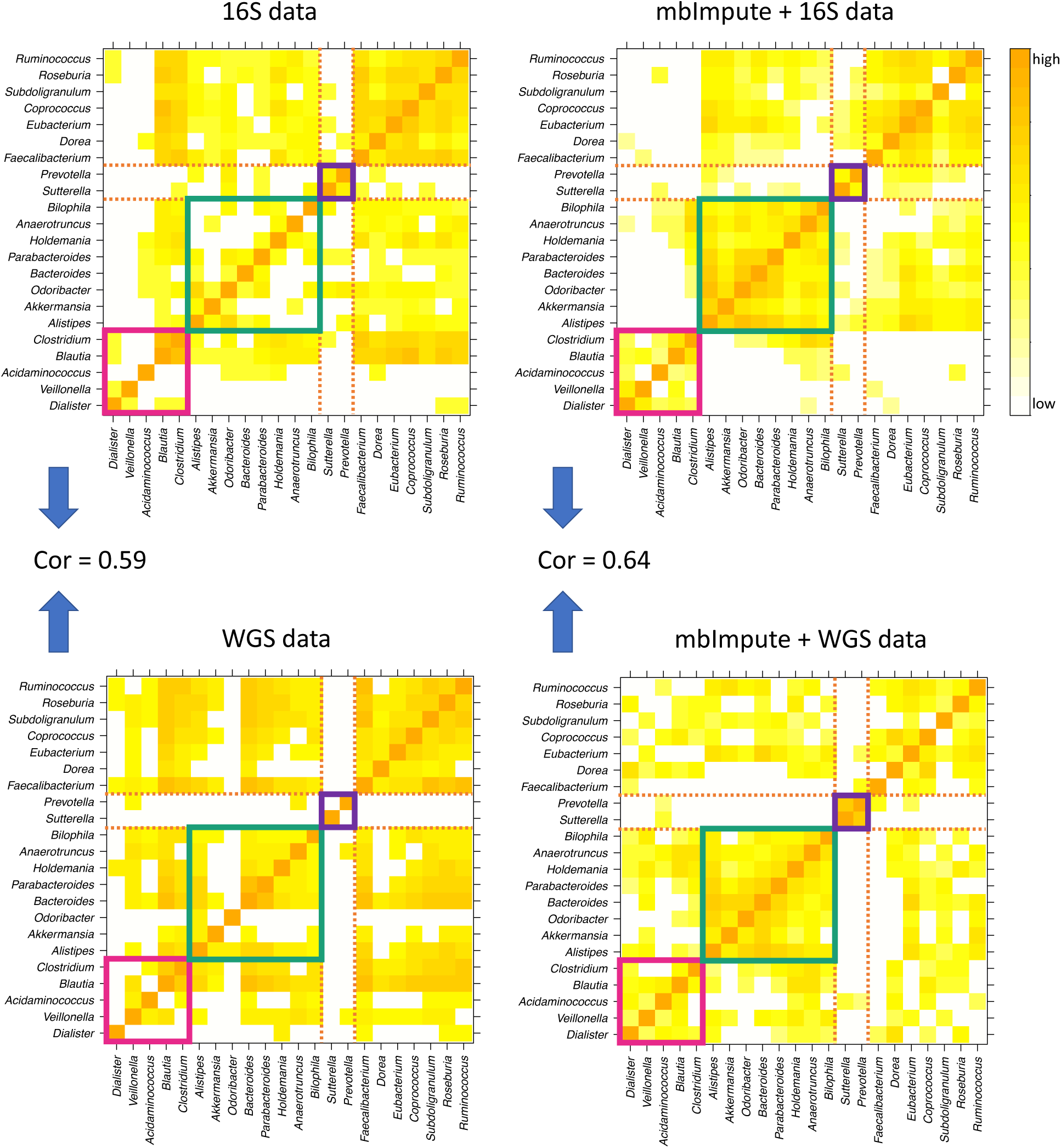
mbImpute improves the consistency in estimating taxon-taxon correlations between 16S and WGS data of microbiome composition in the healthy human stool samples. Four Pearson correlation matrices are calculated based on genus-level taxa’s abundances in 16S and WGS data, with or without using mbImpute as a preceding step. Before imputation, the Pearson correlation between the two correlation matrices is 0.59, and this correlation increases to 0.64 after imputation. For illustration purposes, each heatmap shows square roots of Pearson correlations, with the bottom 40% of values truncated to 0. The magenta, green, and purple squares highlight three taxon groups, each of which contains strongly correlated taxa and is consistent between the 16S and WGS data after imputation.

## Discussion

A critical challenge in microbiome data analysis is statistical inference of taxon abundance from highly sparse and noisy data. Our proposed method, mbImpute, will address this challenge and facilitate analysis of both 16S and WGS data. mbImpute works by correcting non-biological zeros and retaining taxa’s non-zero abundance distributions after imputation. As the first imputation method designed for microbiome data, mbImpute is shown to outperform multiple state-of-the-art imputation methods developed for other data types. Regarding applications of mbImpute, we demonstrate that the mbImpute-empowered DA analysis has advantages over the existing DA methods in three aspects. First, mbImpute increases the power of DA taxon identification by recovering the taxa that are missed by the existing methods (due to excess zeros) but should be called DA (i.e., having non-zero abundances exhibiting different means between two sample groups). Second, mbImpute reduces the false positive taxa, which are identified by the existing methods (due to different proportions of zeros) but should not be called DA (i.e., having similar non-zero abundances between two sample groups). Third, mbImpute improves the reproducibility of DA taxon identification across studies and the consistency of microbial community detection between 16S and WGS data. Furthermore, we found literature evidence for the DA taxa identified as enriched in T2D or CRC samples after mbImpute was applied, supporting the application potential of mbImpute in revealing microbiome markers for disease diagnosis and therapeutics.

There has been a long-standing concern about sample contamination in microbiome sequencing data, e.g. contamination from DNA extraction kits and laboratory reagents [1, 3]. Existing studies have attempted to address this issue via calibrated sequencing operations [2, 3, 6] and computational methods [4,5]. We recommend researchers to perform contamination removal before applying mbImpute. Moreover, by its design, mbImpute is robust to certain types of sample contamination that result in outlier taxa and samples. For each outlier taxon, mbImpute would borrow little information from other taxa to impute this outlier taxon’s abundances. Similarly, mbImpute is robust to the existence of outlier samples that do not resemble any other sample. In statistical inference, a popular and powerful technique is the use of indirect evidence by borrowing information from other observations, as seen in regression, shrinkage estimation, empirical Bayes, among many others [93]. Imputation follows the indirect evidence principle, where the most critical issue is to decide what observations to borrow information from so as to improve data quality instead of introducing excess biases. To achieve this, mbImpute employs penalized regression to selectively leverage similar samples, similar taxa, and sample covariates to impute likely non-biological zeros, whose identification also follows the indirect evidence principle by incorporating sample covariates into consideration. mbImpute also provides a flexible framework to make use of microbiome metadata: it selectively borrows metadata information when available, but it does not rely on the existence of metadata (see Methods).

In the comparison of mbImpute with softImpute, a general matrix imputation method widely used in other fields, we observed that softImpute’s imputed taxon abundances exhibit artificial spikes and smaller variances than those of the original non-zero abundances, possibly due to its low-rank assumption. In contrast, mbImpute is a regression-based method that focuses more on local matrix structures, and we found that it retains well the original non-zero abundance distributions. We will investigate the methodological differences between mbImpute and softImpute in a future study.

Moreover, we observed that, similar to each taxon’s non-zero abundances, the imputed abundances exhibit a bell-shaped distribution across samples on the logarithmic scale. This suggests that statistical methods utilizing Normal distributional assumptions become suitable and applicable to imputed taxon abundances. For example, we have shown that the two-sample *t*-test works well with the imputed data in the identification of DA taxa. In addition to DA analysis, another possible use of the imputed microbiome data is to construct a taxon-taxon interaction network, to which network analysis methods may be applied to find taxon modules and hub taxa [94]. As a preliminary exploration, we constructed Bayesian networks of taxa based on the two T2D datasets Qin et al. and Karlsson et al. after applying mbImpute. Interesting shifts are observed in taxon interactions from control samples to T2D samples (Supplementary Figs. S7–8). For example, two genera, *Ruminococcus* and *Eubacterium*, have interactive species in control samples but not in T2D samples. In future research, differential network analysis methods can be applied to find taxon communities whose interactions differ between two sample groups.

## Methods

### mbImpute methodology

Here we describe mbImpute, a statistical method that corrects prevalent non-biological zeros in microbiome data. As an overview, mbImpute takes an taxon count matrix as input, pre-processes the data, identifies the likely non-biological zeros and imputes them based on the input count matrix, sample metadata, and taxon phylogeny, and finally outputs an imputed count matrix.

#### Notations

We denote the sample-by-taxon taxa count matrix as 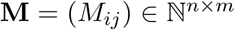, where *n* is the number of samples and *m* is the number of taxa. We denote the sample covariate matrix (i.e., metadata) as **X** ∈ ℝ^*n×q*^, where *q* denotes the number of covariates plus one (for the intercept). (By default, mbImpute includes sample library size as a covariate.) In addition, we define a phylogenetic distance matrix of taxa as 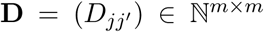, where *D*_*jj′*_ represents the number of edges connecting taxa *j* and *j*^′^ in the phylogenetic tree.

#### Data pre-processing

mbImpute requires every taxon’s counts across samples to be on the same scale before imputation. If this condition is unmet, normalization is needed. However, how to properly normalize microbiome data is challenging, and multiple normalization methods have been developed in recent years [29, 95, 96]. To give users the flexibility of choosing an appropriate normalization method, mbImpute allows users to directly input a normalized count matrix by specifying that the input matrix does not need normalization. Otherwise, mbImpute normalizes samples by library size.

###### Default normalization (optional)

To account for the varying library sizes (i.e., total counts) of samples, mbImpute first normalizes the count matrix M by row. The normalized count matrix is denoted as 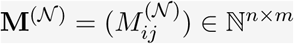, where

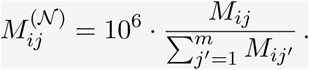

After this normalization, every sample has a total count of 10^6^.

First, mbImpute filters out taxa that have too few non-zero counts to avoid imputing these taxa’s zeros, which are likely biological. This filtering step is exactly the same as how Kaul et al. [30] define structural zeros, i.e., true zeros. More specifically, taxon *j* would be filtered out if the 95% confidence interval of its expected non-zero proportion does not cover zero:

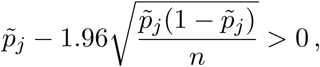

where 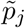 is the observed non-zero proportion of taxon *j*. This filtering step is called the binomial test. In the mbImpute package, users can choose the filtering threshold.

Next, mbImpute applies the logarithmic transformation to the normalized counts so as to reduce the effects of extremely large counts [97]. The resulted log-transformed normalized matrix is denoted as 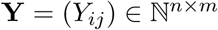, with

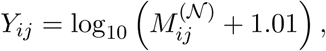

where the value 1.01 is added to make *Y*_*ij*_ > 0 to avoid the occurrence of infinite values in a later parameter estimation step, following Li and Li [50, 98]. This logarithmic transformation allows us to fit a continuous probability distribution to the transformed data, thus simplifying the statistical modeling. In the following text, we refer to **Y** as the sample-by-taxon abundance matrix.

#### mbImpute step 1: identification of taxon abundances that need imputation

mbImpute assumes that each taxon’s abundances (across samples within a sample group), i.e., a column in **Y**, follow a mixture model. The model consists of two components: a Gamma distribution for the taxon’s false zero and low abundances and a Normal distribution for the taxon’s actual abundances, with the Normal mean incorporating sample covariate information (including sample library size as a covariate). Specifically, mbImpute assumes that the abundance of taxon *j* in sample *i*, *Y*_*ij*_, follows the following mixture distribution:

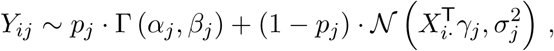

where *p*_*j*_ ∈ (0, 1) is the missing rate, i.e., the probability that taxon *j* is falsely undetected, Γ (*α*_*j*_, *β*_*j*_) denotes the Gamma distribution with shape parameter *α*_*j*_ > 0 and rate parameter *β*_*j*_ > 0, and 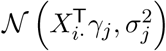 denotes the Normal distribution with mean 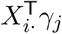 and standard deviation *σ*_*j*_ > 0. In other words, with probability *p*_*j*_, *Y*_*ij*_ is a missing value that needs imputation; with probability 1 − *p*_*j*_, *Y*_*ij*_ does not need imputation but reflects the actual abundance of taxon *j* in sample *i*. mbImpute models the Normal mean parameter as a linear function of sample covariates: 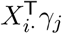, where *X*_*i⋅*_ ∈ ℝ^*q*^ denotes the *i*-th row in the covariate matrix **X**, i.e., the covariates of sample *i*, and *γ*_*j*_ ∈ ℝ^*q*^ represents the *q*-dimensional covariate effect vector of taxon *j*. This allows a taxon to have similar expected (actual) abundances in samples with similar covariates.

The intuition behind this model is that taxon *j’*s actual abundance in a sample (i.e., subject) is drawn from a Normal distribution, whose mean depicts the expected abundance given the sample covariates. However, due to the under-sampling issue in sequencing, false zero or low counts could have been introduced into the data, creating another mode near zero in taxon *j’*s abundance distribution. mbImpute models that mode using a Gamma distribution with mean *α*_*j*_/*β*_*j*_, which is close to zero.

mbImpute fits this mixture model to taxon *j’*s abundances using the Expectation-Maximization (EM) algorithm to obtain the maximum likelihood estimates 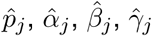, and 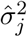. Supplementary Fig. S9 shows four examples where the fitted mixture model well captures the bimodality of an individual taxon’s abundance distribution. However, some taxa are observed to have an abundance distribution containing a single mode that can be well modelled by a Normal distribution. When that occurs, the EM algorithm would encounter a convergence issue. To fix this, mbImpute uses a likelihood ratio test (LRT) to first decide if the Gamma-Normal mixture model fits to taxon *j’*s abundances significantly better than a Normal distribution 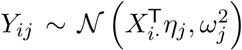 does. Given the maximum likelihood estimates 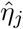 and 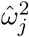 and under the assumption that *Y*_*ij*_’s are all independent, the LRT statistic of taxon *j* is:

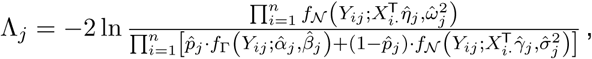

which asymptotically follows a Chi-square distribution with 3 degrees of freedom (because the mixture model has three more parameters than in the Normal model) under the null hypothesis that the Normal model is the correct model. If the LRT p-value ≤ 0.05, mbImpute uses the mixture model to decide which of taxon *j’*s abundances need imputation. Specifically, mbImpute decides if *Y*_*ij*_ needs imputation based on the estimated posterior probability that *Y*_*ij*_ comes from the Gamma component:

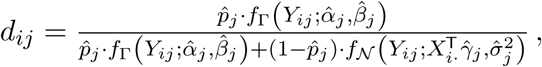

where 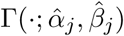 and 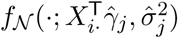 represent the probability density functions of the estimated Gamma and Normal components in the mixture model. Otherwise, if the LRT p-value > 0.05, mbImpute concludes that none of taxon *j’*s abundances need imputation and sets *d*_1*j*_ = … = *d*_*nj*_ = 0.

Based on the *d*_*ij*’_s, mbImpute defines a set Ω of (sample, taxon) pairs whose abundances are unlikely missing and thus do not need imputation:

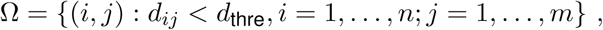

and a complement set Ω^*c*^ containing other (sample, taxon) pairs whose abundances need imputation:

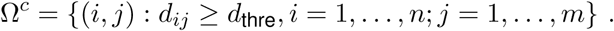

Although *d*_thre_ = 0.5 is used as the default threshold on *d*_*ij*_’s to decide the abundances that need imputation, mbImpute is fairly robust to this threshold choice because most *d*_*ij*_’s are concentrated around 0 or 1. We show this phenomenon in Supplementary Fig. S10, which displays the distribution of all the *d*_*ij*_’s in the data from Zeller et al. [14], Feng et al. [15], Yu et al. [16], Vogtmann et al. [17], Qin et al. [19], and Karlsson et al. [18].

To summarize, mbImpute does not impute all zeros in the taxon count matrix; instead, it first identifies the abundances that are likely missing using a mixture-modelling approach, and it then only imputes these values in the next step.

#### mbImpute step 2: imputation of the missing taxon abundances

In step 1, mbImpute identifies a set Ω of the (sample, taxon) pairs whose abundances do not need imputation. To impute the abundances in Ω^*c*^, mbImpute first learns inter-sample and inter-taxon relationships from Ω by training a predictive model for *Y*_*ij*_, the abundance of taxon *j* in sample *i*. The rationale is that taxon *j* should have similar abundances in similar samples, and that in every sample, the taxa similar to taxon *j* should have abundances similar to taxon *j’*s abundance. In addition, sample covariates are assumed to carry predictive information of taxon abundances. Hence, for interpretability and stability reasons, mbImpute uses a linear model to combine the predictive power of similar taxa, similar samples, and sample covariates:

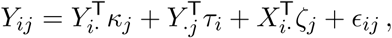

where *Y*_*i*⋅_ ∈ ℝ^*m*^ denotes the *m* taxa’s abundances in sample *i*, *Y*_⋅*j*_ ∈ ℝ^*n*^ enotes taxon *j’*s abundances in the *n* samples, *X*_*i⋅*_ ∈ ℝ^*q*^ denotes sample *i’*s covariates (including the intercept), and *ϵ*_*ij*_ is the error term. The parameters to be estimated include *κ*_*j*_ ∈ ℝ^*m*^, *τ*_*i*_ ∈ ℝ^*n*^ and *ζ*_*j*_ ∈ ℝ^*q*^, *i* = 1, …, *n*; *j* = 1, …, *m*. Note that *κ*_*j*_ represents the *m* taxa’s coefficients (i.e., weights) for predicting taxon *j*, with the *j*-th entry set to zero, so that taxon *j* would not predict itself; *τ*_*i*_ represents the *n* samples’ coefficients (i.e., weights) for predicting sample *i*, with the *i*-th entry set to zero, so that sample *i* would not predict itself; *ζ*_*j*_ represents the coefficients of sample covariates for predicting taxon *j*. In the model, the first term 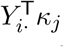 borrows information across taxa, the second term 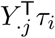 borrows information across samples, and the third term 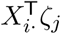 borrows information from sample covariates. The total number of unknown parameters is *m*(*m* − 1)+ *n*(*n* − 1) + *mq*, while our data **Y** and **X** together have *nm* + *nq* values only. Given that often *m » n*, the parameter estimation problem is high dimensional, as the number of parameters far exceeds the number of data points. mbImpute performs regularized parameter estimation by using the Lasso-type *ℓ*_1_ penalty, which leads to good prediction and simultaneously selects predictors (i.e., similar samples and similar taxa) to ease interpretation. That is, mbImpute estimates the above parameters by minimizing the following loss function:

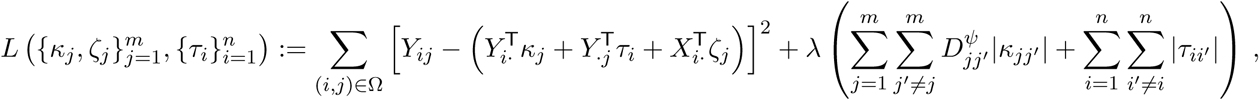

where λ, *ψ* ≥ 0 are tuning parameters chosen by cross-validation, *D*_*jj′*_ represents the phylogenetic distance between taxa *j* and *j*′, *κ*_*jj*_′ represents the *j*′-th element of *κ*_*j*_, and *τ*_*ii*_′ represents the *i*^j^-th element of *τ*_*i*_. Here 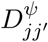, i.e., *D*_*jj′*_ to the power of *ψ*, represents the penalty weight of |*κ*_*jj*_′|.The intuition is that if two taxa are closer in the phylogenetic tree, they are more closely related in evolution and tend to have more similar DNA sequences and biological functions [99, 100], and thus we want to borrow more information between them. For example, if *D*_*j1j2*_ > *D*_*j1j3*_, i.e., taxa *j*_1_ and *j*_2_ are farther away than taxa *j*_1_ and *j*_3_ in the phylogenetic tree, then the estimate of *κ*_*j1j2*_ will be more likely shrunk to zero than the estimate of *κ*_*j1j3*_, and mbImpute would use taxon *j*_3’_s abundance more than taxon *j*_2’_s to predict taxon *j*_1’_s abundance. The tuning parameter *ψ* is introduced because the distance *D*_*jj′*_, the number of edges connecting taxa *j* and *j*′, may not be the best penalty weight for prediction purpose. Choosing *ψ* by cross-validation is expected to enhance the predication accuracy.

mbImpute performs the estimation using the `R` package `glmnet` [101] and obtains the parameter estimates: 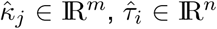, and 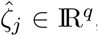, *i* = 1, …, *n*; *j* = 1, …, *m*. Finally, for (*i, j*) ∈ Ω^*c*^, the abundance of taxon *j* in sample *i* is imputed as:

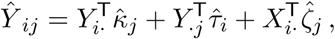

and mbImpute does not alter *Y*_*ij*_ if (*i, j*) ∈ Ω.

Note that mbImpute does not require the availability of the sample covariate matrix **X** or the phylogenetic tree. In the absence of sample covariates, the loss function becomes

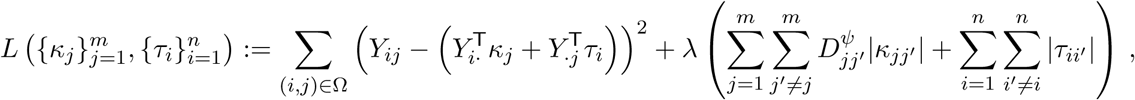

minimizing which returns the parameter estimates: 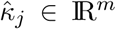 and 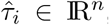, *i* = 1, …, *n*; *j* ≠ {1, …, *m*}. Finally, for (*i, j*) ∈ Ω^*c*^, the abundance of taxon *j* in sample *i* is imputed as:

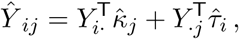

and mbImpute does not alter *Y*_*ij*_ if (*i, j*) ∈ Ω. In the absence of the phylogenetic tree, mbImpute sets *D*_*jj′*_ = 1 for all *j* ≠ *j*′ ∈ {1, …, *m*}.

When *m* is large, mbImpute does not estimate *m*(*m −* 1) + *n*(*n −* 1) + *mq* parameters but uses the following strategy to increase its computational efficiency. For each taxon *j*, mbImpute selects the *k* taxa closest to it (excluding itself) in phylogenetic distance and sets the other (*n − k*) taxa’s coefficients in κ_*j*_ to zero. This strategy reduces the number of parameters to *mk* + *n*(*n −* 1) + *mq* and the computational complexity from *O*(*m*^2^) to *O*(*m*).

In summary, mbImpute step 2 includes two phases: training on Ω and prediction (imputation) on Ω^*c*^, as illustrated in Supplementary Fig. S1.

### Imputation methods

We compared mbImpute with five existing imputation methods designed for non-microbiome data: softImpute and four scRNA-seq imputation methods (scImpute, SAVER, MAGIC, and ALRA). All these imputation methods take a count matrix as input and ouput an imputed count matrix with the same dimensions.

#### softImpute

We used `R` package `softImpute` (version 1.4) and the following command to impute an taxon count matrix (a sample-by-taxon matrix):

~~~
complete(taxa_count_matrix, softImpute(taxa_count_matrix, rank.max = cv.rankmax))
where rank.max was chosen by 10-fold cross-validation.
~~~

#### scImpute

We used `R` package `scImpute` (version 0.0.9) with the input as a taxon-by-sample count matrix (transpose of the matrix in Fig. 1):

~~~
scimpute(count path = “taxa_count_matrix_trans.csv”, Kcluster = 1, out dir = “sim_imp”) where taxa_count_matrix_trans.csv is the input file containing the transposed taxon count matrix.
~~~

### SAVER

We used `R` package `SAVER` (version 1.1.2) with the input as a taxon-by-sample count matrix (transpose of the matrix in Fig. 1):

~~~
saver(t(taxa_count_matrix), ncores = 1, estimates.only = TRUE)
~~~

### MAGIC

We used `Python` package `MAGIC` (version 2.0.3) and the following commands to impute an taxon count matrix:

~~~
magic_op = magic.MAGIC()
magic_op.set params(n_pca = 40)
magic_op.fit_transform(taxa_count_matrix)
~~~

### ALRA

We applied `R` functions `normalize_data`, `choose_k`, and `alra`, which were released on Aug 10, 2019 at https://github.com/KlugerLab/ALRA, and the following commands to impute an taxon count matrix:

~~~
normalized_mat = normalize_data(taxa_count_matrix)
k_chosen = choose_k(normalized_mat, K = 49, noise_start = 44)$k
alra(normalized_mat, k = k_chosen)$A_norm_rank_k_cor_sc
~~~

### DA analysis methods

In both simulation and real data studies, we compared the mbImpute-empowered *t*-test and the softImpute-empowered *t*-test, which apply to log-transformed taxon abundances. We further compared five existing DA methods: the Wilcoxon rank-sum test, ANCOM, ZINB/NB-GLM, metagenomSeq and DESeq2-phyloseq, which apply to taxon counts, with or without using mbImpute as a preceding step. Each method calculates a p-value for each taxon and identifies the DA taxa by setting a p-value threshold to control the false discovery rate (FDR). See Supplementary for the statistical definitions of DA taxa.

#### Wilcoxon rank-sum test

We implemented the Wilcoxon rank-sum test using the `R` function `pairwise.wilcox.test` in the package `stats` (version 3.5.1). For each taxon, we performed the test on its counts in two sample groups to obtain a p-value, which suggests if this taxon is DA between the two groups. In simulations, we used the following command to implement a two-sided test for each taxon: `pairwise.wilcox.test(x = taxon counts, g = condition, p.adjust.method = “none”)` In real data analysis, we used the following command to implement a one-sided test to find if a taxon is disease-enriched (the first condition is the disease condition) and obtained a p-value: `pairwise.wilcox.test(x = taxon counts, g = condition, p.adjust.method = “none”, alternative = “greater”)`

### ANCOM

We used the `ANCOM.main` function released on Sep 27, 2019 at https://github.com/FrederickHuangLin/ `ANCOM` [27]. Since this function does not provide an option for a one-sided test, we used its default settings and reported its identified DA taxa based on a two-sided test with a significance level 0.1 (`sig = 0.1`), in both simulations and real data analysis. We note that no external FDR control was implemented. Specifically, we used the following command to obtain the result of ANCOM: `ANCOM.main(taxa count matrix, covariate matrix, adjusted = F, repeated = F, main.var = “condition”, adj.formula = NULL, repeat.var = NULL, multcorr = 2, sig = 0.1, prev.cut = 0.90, longitudinal = F)` where `taxa count matrix` is a sample-by-taxon count matrix and `covariate_matrix` is a sample-by-covariate matrix, same as the input of mbImpute.

### ZINB-GLM

We implemented the ZINB-GLM method using the `R` function `zeroinfl` in the package `pscl` (version 1.5.2). For each taxon, the `condition` variable is a group indicator (treatment or control) included as a predictor in the generalized linear model (GLM). The partial Wald test was used to test if the coefficient of the `condition` variable is significantly different from 0. For each taxon, we used the following command to implement the ZINB-GLM method:

~~~
summary(zinb <- zeroinfl(taxa count matrix[,i] ∼ condition, dist = “negbin”))
~~~

In simulations, we used the output two-sided p-value for each taxon. In real data analysis, we were interested in the disease-enriched taxa, so we converted the output two-sided p-value into a one-side p-value as follows:

- If the estimated coefficient is non-negative, we divided the p-value by two;
- otherwise, we set the p-value to 1.

#### metagenomeSeq

We used two `R` packages, `metagenomeSeq` combined with `phyloseq`. Specifically, we used the following command to obtain the result:

~~~
mseq obj <- phyloseq to metagenomeSeq(physeq2)
pd <- pData(mseq obj)
mod <- model.matrix(∼ 1 + condition, data = pd)
ran seq <- fitFeatureModel(mseq obj, mod)
~~~

where `physeq2` is an object created from a count matrix and metadata using the `phyloseq` package.

#### DESeq2-phyloseq

We used the `DESeq2` package combined with `phyloseq`. Specifically, we used the following command to obtain the result of DESeq2:

~~~
Deseq2 obj <- phyloseq to deseq2(physeq2, ∼ condition)
results <- DESeq(Deseq2 obj, test=”Wald”, fitType=”parametric”)
~~~

where `physeq2` is an object created from a count matrix and metadata using the `phyloseq` package.

#### mbImpute-empowered *t*-test and softImpute-empowered *t*-test

For mbImpute-empowered *t*-test, we applied mbImpute (in `R` package `mbImpute`, version 0.0.1) to samples in each sample group and then collected the sample groups together to obtain the imputed data, which have the same dimensions as the original data.

For softImpute-empowered *t*-test, we applied softImpute (in `R` package `softImpute`, version 1.4) to samples in each sample group and then collected the sample groups together to obtain the imputed data, which have the same dimensions as the original data. Specifically, we used the following command to obtain the imputed data for a sample group (condition 1):

~~~
complete(raw data condition1, softImpute(raw data condition1, rank.max = cv.rankmax))
~~~

where `rank.max` was chosen by 10-fold cross-validation.

Then for each taxon, we performed the two-sample *t*-test on the imputed data of the scale

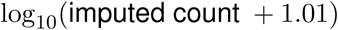

instead of the original count matrix to obtain a p-value, which suggests if this taxon is DA between the two groups. In simulations, we used the following command to implement a two-sided test for each taxon:

~~~
pairwise.t.test(x = taxon imputed, g = condition, p.adjust.method = “none”)
~~~

In real data analysis, we used the following command to implement a one-sided test to find if a taxon is disease-enriched (the first condition is the disease condition) and obtained a p-value:

~~~
pairwise.t.test(x = taxon imputed, g = condition, p.adjust.method = “none”, alternative = “greater”)
~~~

For the Wilcoxon rank-sum test, ZINB-GLM, and mbImpute-empowered or softImpute-empowered *t*-test, after obtaining the p-values of all taxa and collecting them into a vector `p_values`, we adjusted them for FDR control using the `R` function `p.adjust` in the package `stats` (version 3.5.1): `p.adjust`(`p_values`, `method = “fdr”`)

Then we set the FDR threshold to 0.1 in both simulation and real data analysis. The taxa whose adjusted p-values did not exceed this threshold were called DA. ANCOM directly outputs the DA taxa. DESeq2-phyloseq uses the Benjamini-Hochberg procedure to control the FDR under 0.1. For metagenomeSeq, we thresholded its FDR adjusted p-values at 0.1.

### T2D and CRC datasets

We applied mbImpute to six real microbiome datasets, each corresponding to an independent study on the relationship between microbiomes and the occurrence of a human disease. All these six datasets were generated by the whole genome shotgun sequencing and are available in the `R` package `curatedMetagenomicData` [102]. We compared the disease-enriched DA taxa identified by each of four DA methods, namely the Wilcoxon rank-sum test, ANCOM, ZINB-GLM, and the mbImpute-empowered *t*-test. Below is the description of the six datasets and our analysis.

Two datasets are regarding T2D [18, 19]. The Karlsson *et al.* data contain 145 fecal samples from 70-year-old European women for studying the relationship between human gut microbiome compositions and T2D status. The samples/subjects are in three groups: 53 women with T2D, 49 women with impaired glucose tolerance (IGT), and 43 women as the normal control (CON). The twelve sample covariates include the study condition, the subject’s age, the number of reads in each sample, the triglycerides level, the hba1c level, the ldl (low-density lipoprotein cholesterol) level, the c peptide level, the cholesterol level, the glucose level, the adiponectin level, the hscrp level, and the leptin level. In our analysis, we considered the 344 taxa at the species level with phylogenetic information available in the `R` package `curatedMetagenomicData`. Qin et al. [19] performed deep shotgun metagenomic sequencing on 369 Chinese T2D patients and non-diabetic controls (CON). The three sample covariates include the study condition, the body mass index, and the number of reads in each sample. We analyzed 469 taxa at the species level with phylogenetic information. From both datasets, we identified T2D-enriched taxa by comparing the T2D and CON groups.

Four datasets are regarding CRC –17]. Zeller et al. [14] and Feng et al. [15] studied CRC-related microbiomes in three conditions: CRC, small adenoma (ADE; diameter < 10 mm), and control (CON). Zeller et al. [14] sequenced the fecal samples of patients across two countries (France and Germany) in these three groups: 191 patients with CRC, 66 patients with ADE, and 42 patients in CON. The sample covariates include the study condition, the subject’s age category, gender, body mass index and country, and the number of reads in each sample. We included 486 taxa at the species level with phylogenetic information. Feng et al. [15] sequenced samples from 154 human subjects aged between 45–86 years old in Australia, including 46 patients with CRC, 47 patients with ADE, and 61 in CON. The sample covariates include the study condition, the subject’s age category, gender and body mass index, and number of reads in each sample. We included 449 taxa at the species level in our analysis. Yu et al. [16] and Vogtmann et al. [17] studied CRC-related microbiomes in two conditions: CRC vs. CON. In detail, Yu et al. [16] sequenced 128 Chinese samples, including 75 patients with CRC and 53 patients in CON. The sample covariates include the study condition and the number of reads in each sample. We studied 417 taxa at the species level. Vogtmann et al. [17] included 104 samples from Washington DC and sequenced their fecal samples, including 52 with CRC and 52 in CON. The sample covariates include the study condition, the subject’s age category, gender and body mass index, and number of reads in each sample. We included 412 taxa at the species level. From all the four datasets, we identified CRC-enriched taxa by comparing the CRC and CON groups.

## Software and code

The `mbImpute R` package and the code for simulation and real data analysis are available at https://github.com/ruochenj/mbImpute

## Acknowledgements

The authors would like to thank Dr. Hongzhe Li at University of Pennsylvania for pointing us to this research direction. The authors also appreciate the comments and feedback from the members of the Junction of Statistics and Biology at UCLA (http://jsb.ucla.edu).

## Funding

This work was supported by the following grants: National Science Foundation DMS-1613338 and DBI-1846216, NIH/NIGMS R01GM120507, PhRMA Foundation Research Starter Grant in Informatics, Johnson & Johnson WiSTEM2D Award, and Sloan Research Fellowship (to J.J.L).

## Notes

### Competing Interest Statement

The authors have declared no competing interest.

### Summary of Updates

In summary, we have made the following six major changes in our revised manuscript, and these changes are summarized in Figures 2, 3, 5, and 6 and Supplementary Figures S3, S4, S7, and S8. I. We show that mbImpute works for both 16S rRNA sequencing and whole-genome sequencing (WGS) microbiome data. II. We have performed more comprehensive analyses to show that mbImpute can improve the accuracy of multiple existing methods for identifying differentially abundant (DA) taxa from microbiome data;. First, we have added two more DA methods, metagenomeSeq and DEseq2-phyloseq, to both simulation and real data DA studies. This addition makes our analysis include six DA methods, and we show that all these methods exhibit improved accuracy after mbImpute is applied. Second, we have added sparseDOSSA, a 16S data simulator, to generate 16S data, and we show that mbImpute is effective in improving the accuracy of all the six DA methods on the 16S data. III. We have added a new accuracy criterion for the DA taxon analysis on real data: the accuracy of using DA taxa to predict sample conditions across studies. In detail, our cross-study results show that, after mbImpute, the DA taxa identified from the data of one study become more predictive of the sample conditions (e.g., diseased vs. undiseased) of another study of the same disease. IV. We have added a new result to show that mbImpute preserves distributional characteristics of taxa's non-zero abundances and recovers downsampling. Our new analysis shows that pairwise taxon-taxon relationships inferred from non-zero abundances can be preserved by mbImpute. In addition, we show that mbImpute can effectively recover zeros introduced by downsampling. V. We have added a concordance analysis between 16S data and WGS data after imputation. Specifically, we have compared the genus correlations in 16S data and WGS data before and after imputation. We observe that, before imputation, the genus correlations are vastly different between 16S data and WGS data. After mbImpute, the genus correlations become more similar between the two data types. Based on literature, we have further verified the highly correlated genera found in both types of data after imputation. VI. We have added preliminary results of using mbImpute to assist the construction of taxon interaction networks.

