## Supplementary File for "mbImpute: an accurate and robust imputation method for microbiome data"

#### Simulation 1 for benchmarking imputation methods

To compare mbImpute with existing imputation methods developed for non-microbiome data, we generated microbiome abundances from a generative model fitted to the T2D data with $53$ subjects and $344$ taxa (Karlsson *et al.*, 2013). Below we describe the data generation process step by step.

##### Complete data generation

1. We followed the data pre-processing steps of mbImpute (see Methods in the main text) to convert the OTU count matrix to log-transformed normalized abundances. Then we removed the taxa with greater than $95\%$ of zero counts (equivalently, $\log_{10}(1.01)$ abundances) across subjects and kept $193$ taxa. We denote the abundance matrix after this filtering step by $\mathbf{Y} = (Y_{ij}) \in \mathbb{R}^{n \times m}$, where $n = 53$ and $m = 193$. We also collected the sample covariate matrix $\mathbf{X} \in \mathbb{R}^{n \times q}$, where $q = 12$.
2. Following step 1 of mbImpute (see Methods in the main text), we identified a set $\Omega$ of (sample, taxon) pairs whose abundances are unlikely missing and thus do not need imputation.
3. Following step 2 of mbImpute (see Methods in the main text), we fit the following model to the (sample, taxon) pairs in $\Omega$:

$$Y_{ij} = Y_{i\cdot}^{\mathsf{T}} \kappa_j + Y_{\cdot j}^{\mathsf{T}} \tau_i + X_{i\cdot}^{\mathsf{T}} \zeta_j + \epsilon_{ij} \, ,$$

[1] Department of Statistics, University of California, Los Angeles, CA 90095-1554
[2] Department of Biostatistics and Epidemiology, Rutgers School of Public Health, Rutgers, New Jersey 08854
[3] Department of Human Genetics, University of California, Los Angeles, CA 90095-7088
[4] Department of Computational Medicine, University of California, Los Angeles, CA 90095-1766

by minimizing the loss function

$$\sum_{(i,j)\in\Omega}\left(Y_{ij}-\left(Y_{i\cdot}^{\mathsf{T}}\kappa_j+Y_{\cdot j}^{\mathsf{T}}\tau_i+X_{i\cdot}^{\mathsf{T}}\zeta_j\right)\right)^2+\lambda\left(\sum_{j=1}^{m}\sum_{j'\neq j}^{m}D_{jj'}^{\psi}|\kappa_{jj'}|+\sum_{i=1}^{n}\sum_{i'\neq i}^{n}|\tau_{ii'}|\right)$$

and obtaining the parameter estimates: $\hat{\kappa}_j\in\mathbb{R}^m$, $\hat{\tau}_i\in\mathbb{R}^n$, and $\hat{\zeta}_j\in\mathbb{R}^q$, $i=1,\ldots,n$; $j=1,\ldots,m$. The tuning parameters $\lambda,\psi\geq 0$ were chosen by cross-validation. Note that we constrained $\kappa_j$ to have only $5$ non-zero entries corresponding to the $5$ taxa closest to taxon $j$ in phylogenetic distance, i.e., $\{j' : D_{jj'}$ is among the five smallest of all $D_{jr}, r\neq j\}$.

4. We generated the complete, log-transformed abundance of taxon $j$ in sample $i$, as

$$Y_{ij}^{\text{comp}}=Y_{i\cdot}^{\mathsf{T}}\hat{\kappa}_j+Y_{\cdot j}^{\mathsf{T}}\hat{\tau}_i+X_{i\cdot}^{\mathsf{T}}\hat{\zeta}_j\,.$$

We denote the resulting matrix $\mathbf{Y}^{\text{comp}}=(Y_{ij}^{\text{comp}})$ as the complete data that contain log-transformed abundances without missing values.

#### Zero-inflated data generation

Next, we introduced zero inflation to $\mathbf{Y}^{\text{comp}}$ and generated $\mathbf{Y}^{\text{zi}}$ by mimicking real data as follows. With the identified $\Omega$, we calculated $z_k^{\text{real}}$ (taxon $k$'s proportion of likely false zeros across samples) and $\mu_k^{\text{real}}$ (taxon $k$'s average abundance after we excluded likely false zeros) for each taxon $k$ in Karlsson *et al.*'s data $\mathbf{Y}$, $k=1,\ldots,m$. Next we introduced zeros into $\mathbf{Y}^{\text{comp}}$ in the following non-parametric way for each taxon $j$, $j=1,\ldots,m$:

1. We calculated taxon $j$'s average abundance in $\mathbf{Y}^{\text{comp}}$ as the mean of column $j$, denoted by $\mu_j^{\text{comp}}$;
2. We randomly sampled a value from $\{z_k^{\text{real}} : \mu_k^{\text{real}}\in(\mu_j^{\text{comp}}-0.5,\mu_j^{\text{comp}}+0.5)\}$ and denoted it as $z_j^{\text{zi}}$, i.e., the proportion of false zeros to be introduced into taxon $j$'s abundances in $\mathbf{Y}^{\text{comp}}$;
3. We randomly drew false zero indicators $I_{ij}\sim$ Bernoulli$(z_j^{\text{zi}})$ independently for sample $i=1,\ldots,n$;
4. We set $Y_{ij}^{\text{zi}}=\max\left(Y_{ij}^{\text{comp}}\cdot I_{ij},\,\log_{10}(1.01)\right)$.

The reason why we set the minimum value of $\mathbf{Y}^{\text{zi}}$ to $\log_{10}(1.01)$ is that mbImpute step 1 sets the minimum log-transformed abundance to $\log_{10}(1.01)$ to facilitate the fitting of the Gamma-Normal mixture model.

#### Evaluation criteria for imputation accuracy

After applying an imputation method to $\mathbf{Y}^{\text{zi}}$ (mbImpute also used $\mathbf{X}$), we obtained $\mathbf{Y}^{\text{imp}}$ and evaluated the imputation performance by the following three criteria. Note that mbImpute identified $8$ taxa with too many zeros in $\mathbf{Y}^{\text{zi}}$ and excluded them from imputation. For a fair comparison, we excluded these $8$ taxa from the calculation of the evaluation criteria for every imputation method, so $m$ was reduced to $193 - 8 = 185$ in the following.

1. Mean squared error (MSE) between $\mathbf{Y}^{\text{imp}}$ and $\mathbf{Y}^{\text{comp}}$:

$$\text{MSE} = \frac{1}{nm} \sum_{i=1}^{n} \sum_{j=1}^{m} (Y_{ij}^{\text{comp}} - Y_{ij}^{\text{imp}})^2 ,$$

   which is shown in Fig. 2a.
2. Pearson correlation between $Y_{\cdot j}^{\text{imp}}$ and $Y_{\cdot j}^{\text{comp}}$ for $j = 1, \ldots, m$. The mean of these $m$ correlations is shown in Fig. 2b.
3. Mean and standard deviation (SD) of taxon $j$ in the imputed data vs. those in the complete data:

$$\bar{Y}_{\cdot j}^{\text{imp}} = \frac{1}{n} \sum_{i=1}^{n} Y_{ij}^{\text{imp}} \quad \text{vs.} \quad \bar{Y}_{\cdot j}^{\text{comp}} = \frac{1}{n} \sum_{i=1}^{n} Y_{ij}^{\text{comp}} ,$$

$$\text{sd}_{\cdot j}^{\text{imp}} = \sqrt{\frac{1}{n-1} \sum_{i=1}^{n} \left( Y_{ij}^{\text{imp}} - \bar{Y}_{\cdot j}^{\text{imp}} \right)^2} \quad \text{vs.} \quad \text{sd}_{\cdot j}^{\text{comp}} = \sqrt{\frac{1}{n-1} \sum_{i=1}^{n} \left( Y_{ij}^{\text{comp}} - \bar{Y}_{\cdot j}^{\text{comp}} \right)^2} ,$$

   $j = 1, \ldots, m$. In Fig. 2c, we computed the Wasserstein distance between the distribution of $\left\{ \bar{Y}_{\cdot 1}^{\text{imp}} / \text{sd}_{\cdot 1}^{\text{imp}}, \ldots, \bar{Y}_{\cdot m}^{\text{imp}} / \text{sd}_{\cdot m}^{\text{imp}} \right\}$ vs. that of $\{ \bar{Y}_{\cdot 1}^{\text{comp}} / \text{sd}_{\cdot 1}^{\text{comp}}, \ldots, \bar{Y}_{\cdot m}^{\text{comp}} / \text{sd}_{\cdot m}^{\text{comp}} \}$. In Fig. 2d, we computed the Euclidean distance between the imputed data and the complete data as

$$\sqrt{\sum_{j=1}^{m} \left( \bar{Y}_{\cdot j}^{\text{imp}} - \bar{Y}_{\cdot j}^{\text{comp}} \right)^2 + \left( \text{sd}_{\cdot j}^{\text{imp}} - \text{sd}_{\cdot j}^{\text{comp}} \right)^2} .$$

### Simulation 2 for benchmarking imputation methods based on real WGS data

To further benchmark mbImpute against existing imputation methods developed for non-microbiome data, we used a semi-simulation approach by obtaining a subset of microbiome WGS data with at least $86\%$ non-zeros from a T2D dataset composed of $344$ subjects and $469$ taxa (Qin *et al.*, 2012). Below we describe the data generation process step by step.

#### Complete data generation

1. We followed the data pre-processing steps of mbImpute (see Methods in the main text) to convert the OTU count matrix to log-transformed normalized abundances, denoted by $\mathbf{Y} = (Y_{ij}) \in \mathbb{R}^{344 \times 469}$. We also collected the sample covariate matrix $\mathbf{X} \in \mathbb{R}^{344 \times 3}$.
2. Following step 1 of mbImpute (see Methods in the main text), we identified a set $\Omega$ of (sample, taxon) pairs whose abundances are unlikely missing and thus do not need imputation.

3. For each taxon, we checked if it has at least $43$ non-zero counts, or equivalently, at least $43$ abundances greater than $\log_{10}(1.01)$, across the $344$ subjects. If yes, we kept the taxon; otherwise, we filtered it out. This step left us with $145$ taxa.
4. Denote the index set of samples where taxon $j$ has non-zero counts as

$$\mathcal{I}_j = \{i : Y_{ij} > \log_{10}(1.01),\ i = 1, \ldots, 344\}$$

   Note that all $|\mathcal{I}_j| \geq 43$ due the the filtering step. Then we constructed a new index set of samples $\mathcal{I}'_j$, with $|\mathcal{I}'_j| = 50$, in the following way:

   (a) if $|\mathcal{I}_j| < 50$, $\mathcal{I}'_j = \mathcal{I}_j \cup$ a random subset of $\mathcal{I}_j^c$ with size $50 - |\mathcal{I}_j|$;

   (b) otherwise, $\mathcal{I}'_j =$ a random subset of $\mathcal{I}_j$ with size $50$.

5. We constructed the complete data by column-stacking taxon $j$'s abundances in the samples in $\mathcal{I}'_j$, with the $50$ samples randomly ordered, $j = 1, \ldots, 145$, resulting in $\mathbf{Y}^{\text{comp}} \in \mathbb{R}^{50\times 145}$, which we assumed to have no missing values. By our construction, $\mathbf{Y}^{\text{comp}}$ has at least $86\%$ values greater than $\log_{10}(1.01)$, corresponding to non-zero values on the count scale.
6. We constructed the sample covariate matrix $\mathbf{X}^{\text{comp}} \in \mathbb{R}^{50\times 3}$ as follows: for sample $i$ and covariate $r$ ($i = 1, \ldots, 50$; $r = 1, 2, 3$),

   (a) if covariate $r$ is categorical (e.g., gender), $X_{ij}^{\text{comp}}$ was decided by the majority vote of the $j$-th covariate values of the original samples rearranged into the $i$-th row of $\mathbf{Y}^{\text{comp}}$: majority($\{X_{i'j}$ : sample $i'$ is in the $i$-th row of $\mathbf{Y}^{\text{comp}}\}$);

   (b) if covariate $r$ is numerical (e.g., BMI), $X_{ij}^{\text{comp}}$ was set to the average of the $j$-th covariate values of the original samples rearranged into the $i$-th row of $\mathbf{Y}^{\text{comp}}$: average($\{X_{i'j}$ : sample $i'$ is in the $i$-th row of $\mathbf{Y}^{\text{comp}}\}$).

#### Zero-inflated data generation for mimicking WGS data

Next, we introduced zero inflation to $\mathbf{Y}^{\text{comp}}$ and generated $\mathbf{Y}^{\text{zi}}$ by mimicking real WGS data as follows. With the identified $\Omega$, we calculated $z_k^{\text{real}}$ (taxon $k$'s proportion of likely false zeros across samples) and $\mu_k^{\text{real}}$ (taxon $k$'s average abundance after we excluded likely false zeros) for each taxon $k$ in Qin *et al.*'s data, $k = 1, \ldots, 469$. Next we introduced zeros into $\mathbf{Y}^{\text{comp}}$ in the following non-parametric way for each taxon $j$, $j = 1, \ldots, 145$:

1. We calculated taxon $j$'s average abundance in $\mathbf{Y}^{\text{comp}}$ as the mean of column $j$, denoted by $\mu_j^{\text{comp}}$;
2. We randomly sampled a value from $\{z_k^{\text{real}} : \mu_k^{\text{real}} \in (\mu_j^{\text{comp}} - 0.5, \mu_j^{\text{comp}} + 0.5)\}$ and denoted it as $z_j^{\text{zi}}$, i.e., the proportion of false zeros to be introduced into taxon $j$'s abundances in $\mathbf{Y}^{\text{comp}}$;
3. We randomly drew false zero indicators $I_{ij} \sim$ Bernoulli($z_j^{\text{zi}}$) independently for sample $i = 1, \ldots, 50$;

4. We set $Y_{ij}^{\text{zi}} = \max\left(Y_{ij}^{\text{comp}} \cdot I_{ij}, \log_{10}(1.01)\right)$.

The reason why we set the minimum value of $\mathbf{Y}^{\text{zi}}$ to $\log_{10}(1.01)$ is that mbImpute step 1 sets the minimum log-transformed abundance to $\log_{10}(1.01)$ to facilitate the fitting of the Gamma-Normal mixture model.

##### Evaluation criteria for imputation accuracy

After applying an imputation method to $\mathbf{Y}^{\text{zi}}$ (mbImpute also used $\mathbf{X}^{\text{comp}}$), we obtained $\mathbf{Y}^{\text{imp}}$ and evaluated the imputation performance by the following three criteria, where $n = 50$ and $m = 145$.

1. Mean squared error (MSE) between $\mathbf{Y}^{\text{imp}}$ and $\mathbf{Y}^{\text{comp}}$:

$$\text{MSE} = \frac{1}{nm} \sum_{i=1}^{n} \sum_{j=1}^{m} (Y_{ij}^{\text{comp}} - Y_{ij}^{\text{imp}})^2 .$$

2. Pearson correlation between $Y_{\cdot j}^{\text{imp}}$ and $Y_{\cdot j}^{\text{comp}}$ for $j = 1, \ldots, m$. The mean of these $m$ correlations is shown in Fig. 2b.

#### Simulation 3 for evaluating the accuracy and robustness of mbImpute

To evaluate the accuracy and robustness of mbImpute, we simulated microbiome abundances based on real data (Zeller *et al.*, 2014) using four schemes. We set the number of samples to $n = 50$ and the number of taxa to $m = 200$. Under each scheme, we first generated the complete data, as described below.

##### Scheme 1: Covariate

1. We randomly sampled (without replacement) $50$ samples' six covariates (gender, age, disease subtype, BMI, country, and number of reads) from Zeller *et al.*'s data. Then we added a $50$-length vector of ones to the sampled covariates to form a sample covariate matrix $\mathbf{X} \in \mathbb{R}^{n \times q}$, with $q = 7$.
2. We randomly sampled (without replacement) $200$ taxa' normalized and log-transformed abundances (see Methods in the main text) from Zeller *et al.*'s data. We fit the Gamma-Normal mixture model (step 1 of mbImpute) to each sampled taxon $j$, and we denote the estimated coefficient vector in the Normal mean as $\hat{\gamma}_j \in \mathbb{R}^q$, $j = 1, \ldots, m$.
3. We simulated the log-transformed abundance of taxon $j$ in sample $i$, $i = 1, \ldots, n$, from the following model:

$$Y_{ij}^{\text{comp}} = X_{i\cdot}^{\mathsf{T}} \hat{\gamma}_j + \epsilon_{ij} ,$$

where $\epsilon_{ij} \sim \mathcal{N}(0, 1)$ independently. We denote the resulting matrix $\mathbf{Y}^{\text{comp}} = (Y_{ij}^{\text{comp}})$ as the complete data that contain log-transformed abundances without missing values.

##### Scheme 2: Sample

1. We randomly divided $n = 50$ samples into $5$ groups. For each sample $i$, $i = 1, \ldots, n$, we denote its group index by $k(i) \in \{1, \ldots, 5\}$.
2. We generated a sample-to-sample distance matrix $\mathbf{D}^{\text{samp}} = (D_{ii'}^{\text{samp}})_{n \times n}$:

$$D_{ii'}^{\text{samp}} = \begin{cases} 2(|k(i) - k(i')| + 1) & \text{if } i \neq i' \\ 0 & \text{otherwise} \end{cases}$$

   so that two samples from closer groups (in terms of group index) had a smaller distance.
3. We converted the distance matrix $\mathbf{D}^{\text{samp}}$ into a sample-to-sample covariance matrix $\mathbf{\Sigma}^{\text{samp}}$ such that two samples having a smaller distance would have a larger covariance

$$\Sigma_{ii'}^{\text{samp}} = \begin{cases} \exp\left\{(-D_{ii'}^{\text{samp}}/6)^{1.3}\right\} & \text{if } i \neq i' \\ \exp\left\{(-D_{ii'}^{\text{samp}}/6)^{1.3}\right\} + 1 & \text{otherwise} \end{cases},$$

   where the constants $6$ and $1.3$ were chosen to make $\mathbf{\Sigma}^{\text{samp}}$ positive definite.
4. We simulated the log-transformed abundances of taxon $j$ in all the $n$ samples, $j = 1, \ldots, m$, from the following model:

$$Y_{\cdot j}^{\text{comp}} \overset{\text{ind}}{\sim} \mathcal{N}(3 \cdot \mathbf{1}_n, \Sigma^{\text{samp}}),$$

   which we collected as columns to form $\mathbf{Y}^{\text{comp}} = (Y_{ij}^{\text{comp}})$, i.e., the complete data containing log-transformed abundances without missing values.

##### Scheme 3: Taxon

1. We randomly divided $m = 200$ taxa into $10$ groups. For each taxon $j$, $j = 1, \ldots, m$, we denote its group index by $r(j) \in \{1, \ldots, 10\}$.
2. We generated a taxon-to-taxon distance matrix $\mathbf{D}^{\text{taxon}} = (D_{jj'}^{\text{taxon}})_{m \times m}$:

$$D_{jj'}^{\text{taxon}} = \begin{cases} 2(|r(j) - r(j')| + 1) & \text{if } j \neq j' \\ 0 & \text{otherwise} \end{cases}$$

   so that two taxa from closer groups (in terms of group index) had a smaller distance.
3. We converted the distance matrix $\mathbf{D}^{\text{taxon}}$ into a taxa-to-taxa covariance matrix $\mathbf{\Sigma}^{\text{taxon}}$ such that two samples having a smaller distance would have a larger covariance

$$\Sigma_{jj'}^{\text{taxon}} = \begin{cases} \exp\left\{(-D_{jj'}^{\text{taxon}}/12)^{1.3}\right\} & \text{if } j \neq j' \\ \exp\left\{(-D_{jj'}^{\text{taxon}}/12)^{1.3}\right\} + 1 & \text{otherwise} \end{cases},$$

   where the constants $12$ and $1.3$ were chosen to make $\mathbf{\Sigma}^{\text{taxon}}$ positive definite.

4. We simulated the log-transformed abundances of sample $i$ in all the $m$ taxa, $i = 1, \ldots, n$, from the following model:
$$Y_{i\cdot}^{\text{comp}} \overset{\text{ind}}{\sim} \mathcal{N}(3 \cdot \mathbf{1}_m, \mathbf{\Sigma}^{\text{taxon}}) ,$$
which we collected as rows to form $\mathbf{Y}^{\text{comp}} = (Y_{ij}^{\text{comp}})$, i.e., the complete data containing log-transformed abundances without missing values.

##### Scheme 4: Taxon + Sample + Covariate

We combined the data generation procedures from the above three schemes. Following the same notations as above, we simulated the log-transformed abundance of taxon $j$ in sample $i$, $i = 1, \ldots, n$, $j = 1, \ldots, m$, as follows:

1. We generated baseline values, following scheme 1:
$$Y_{ij}^{\text{comp1}} = X_{i\cdot}^{\mathsf{T}} \hat{\gamma}_j + \epsilon_{ij}$$
to form $\mathbf{Y}^{\text{comp1}} = (Y_{ij}^{\text{comp1}})$.
2. We introduced a sample correlation structure as in scheme 2:
$$Y_{\cdot j}^{\text{comp2}} \overset{\text{ind}}{\sim} \mathcal{N}(Y_{\cdot j}^{\text{comp1}}, \mathbf{\Sigma}^{\text{samp}}) ,$$
which we collected as columns to form $\mathbf{Y}^{\text{comp2}} = (Y_{ij}^{\text{comp2}})$
3. We introduced a taxon correlation structure as in scheme 3:
$$Y_{i\cdot}^{\text{comp}} \overset{\text{ind}}{\sim} \mathcal{N}(Y_{i\cdot}^{\text{comp2}}, \mathbf{\Sigma}^{\text{taxon}}) ,$$
which we collected as rows to form $\mathbf{Y}^{\text{comp}} = (Y_{ij}^{\text{comp}})$, i.e., the complete data containing log-transformed abundances without missing values.

Next, we introduced zero inflation to $\mathbf{Y}^{\text{comp}}$ and generated $\mathbf{Y}^{\text{zi}}$ by mimicking real data as follows. We applied the step 1 of mbImpute to Zeller *et al.*'s data, which contain $486$ taxa (Zeller *et al.*, 2014); that is, we identified likely false zeros. Then we calculated $z_k^{\text{real}}$ (taxon $k$'s proportion of likely false zeros across samples) and $\mu_k^{\text{real}}$ (taxon $k$'s average abundance after we excluded likely false zeros) for each taxon $k$ in Zeller *et al.*'s data, $k = 1, \ldots, 486$. Next we introduced zeros into $\mathbf{Y}^{\text{comp}}$ in the following non-parametric way for each taxon $j$, $j = 1, \ldots, m$:

1. We calculated taxon $j$'s average abundance in $\mathbf{Y}^{\text{comp}}$ as the mean of column $j$, denoted by $\mu_j^{\text{comp}}$;
2. We randomly sampled a value from $\{z_k^{\text{real}} : \mu_k^{\text{real}} \in (\mu_j^{\text{comp}} - 0.5, \mu_j^{\text{comp}} + 0.5)\}$ and denoted it as $z_j^{\text{zi}}$, i.e., the proportion of false zeros to be introduced into taxon $j$'s abundances in $\mathbf{Y}^{\text{comp}}$;
3. We randomly drew false zero indicators $I_{ij} \sim \text{Bernoulli}(z_j^{\text{zi}})$ independently for sample $i = 1, \ldots, n$;

4. We set $Y_{ij}^{\text{zi}} = \max\left(Y_{ij}^{\text{comp}} \cdot I_{ij},\, \log_{10}(1.01)\right)$.

The reason why we set the minimum value of $\mathbf{Y}^{\text{zi}}$ to $\log_{10}(1.01)$ is that mbImpute step 1 sets the minimum log-transformed abundance to $\log_{10}(1.01)$ to facilitate the fitting of the Gamma-Normal mixture model.

After we applying mbImpute to $\mathbf{Y}^{\text{zi}}$ (mbImpute also used $\mathbf{X}$), we obtained $\mathbf{Y}^{\text{imp}}$ and evaluated the imputation performance by calculating the mean squared error (MSE) between $\mathbf{Y}^{\text{imp}}$ and $\mathbf{Y}^{\text{comp}}$:

$$\text{MSE}^{\text{mbImpute}} = \frac{1}{nm}\sum_{i=1}^{n}\sum_{j=1}^{m}(Y_{ij}^{\text{comp}} - Y_{ij}^{\text{imp}})^2$$

and the MSE between the zero-inflated matrix $\mathbf{Y}^{\text{zi}}$ and $\mathbf{Y}^{\text{comp}}$:

$$\text{MSE}^{\text{no imputation}} = \frac{1}{nm}\sum_{i=1}^{n}\sum_{j=1}^{m}(Y_{ij}^{\text{comp}} - Y_{ij}^{\text{zi}})^2\,.$$

The results are summarized in Fig. 2e in the main text.

#### Simulation 4 for DA analysis on WGS data

We designed a simulation study to test if mbImpute is able to improve the DA analysis through imputation. We generated three matrices: $\mathbf{Y}^{\text{comp}}$ denotes the complete data; $\mathbf{Y}^{\text{zi}}$ denotes the data containing excess zeros (i.e., with zero inflation) introduced to $\mathbf{Y}^{\text{comp}}$; $\mathbf{Y}^{\text{imp}}$ denotes the data resulted from an imputation method applied to $\mathbf{Y}^{\text{zi}}$. All the three matrices contain log-transformed taxon abundances. We assumed that there are two groups of subjects, and we simulated $120$ subjects, $60$ in each group, and $300$ taxa. We set $100$ taxa to have increased mean abundances and another $100$ taxa to have decreased mean abundances in group 1 compared to group 2. Specifically, for each taxon, we first simulated its $60$ abundances in group 2 from $\mathcal{N}(3, 1.5^2)$. If this taxon was chosen to have an increased mean abundance in group 1, we simulated its $60$ abundances from $\mathcal{N}(4, 1.5^2)$; if it was chosen to have a decreased mean abundance in group 1, we simulated its $60$ abundances from $\mathcal{N}(2, 1.5^2)$; otherwise, we simulated its $60$ abundances in group 1 from $\mathcal{N}(3, 1.5^2)$. Note that all the abundances were sampled independently. These simulated abundances were collected into a sample-by-taxon matrix, i.e., $\mathbf{Y}^{\text{comp}}$.

Then we introduced zero inflation to $\mathbf{Y}^{\text{comp}}$ to generate $\mathbf{Y}^{\text{zi}}$ in the same way as in Simulation 3. The resultant $\mathbf{Y}^{\text{comp}}$ and $\mathbf{Y}^{\text{zi}}$ have $7\%$ and $60\%$ of entries below or equal $\log_{10}(1.01)$, respectively. These percentages indicate the proportions of zeros in the original scale of counts.

To make the three existing DA methods (the Wilcoxon test, ANCOM, and ZINB-GLM) applicable to the simulated data $\mathbf{Y}^{\text{zi}}$, we converted the data scale to counts as follows:

$$M_{ij}^{\text{zi}} = \lfloor 10^{Y_{ij}^{\text{zi}}} - 1.01 \rfloor$$

Then we applied the three existing methods to $\mathbf{M}^{\text{zi}} = (M_{ij}^{\text{zi}})$ to identify DA taxa between the two subject groups.

For imputation-empowered DA analysis, we applied mbImpute and softImpute to each subject group in $\mathbf{Y}^{\text{zi}}$ to first impute data, and we denote the imputed data as $\mathbf{Y}^{\text{imp}}$. Then we performed the two-sample $t$ test on $\mathbf{Y}^{\text{imp}}$ to identify DA taxa.

Fig. 2f in the main text illustrates $\mathbf{Y}^{\text{comp}}$, $\mathbf{Y}^{\text{zi}}$, and $\mathbf{Y}^{\text{imp}}$ (imputed by mbImpute).

To evaluate the performance of DA methods, we defined the true DA taxa based on the results of the two-sample $t$ test applied to $\mathbf{Y}^{\text{comp}}$ (at significance level $\alpha = 0.05$). For a fair comparison, we only considered the taxa that were not filtered out by the binomial test in the data pre-processing step of mbImpute, because these taxa have a reasonable number of non-zero abundances to enable a feasible DA analysis. Then we compared the performance of these DA methods in terms of precision, recall, F1 scores, and area under ROC curves (Fig. 2g).

#### Simulation for DA analysis on 16S rRNA data

We used the `R` package `sparseDOSSA` by Ren *et al.* (2016) to simulate the 16S rRNA sequencing data with known DA taxa. Specifically, we used the following command to obtain simulated data:

```
metadata <- matrix(rbinom(n = 100, size = 1, prob = 0.5), nrow = 1, ncol = 100)
simulated_data <- sparseDOSSA(number_features = 150, number_samples = 100,
percent_spiked = 0.3, UserMetadata = metadata)
```

#### Statistical definitions of DA taxa

Here we list three possible statistical definitions of DA taxa. For a given taxon $j$, we denote $M_j^1$ and $M_j^2$ as its (random) counts in two samples from subject groups $1$ and $2$. The first and most straightforward definition of DA is whether the null hypothesis

$$H_0 : \mathbb{E}[M_j^1] = \mathbb{E}[M_j^2]$$

is rejected. The Wilcoxon rank-sum test is based on this definition. However, this definition has an obvious drawback: it ignores the existence of excess false zeros, i.e., many observed zero values of $M_j^1$ and $M_j^2$ are not reliable. This drawback motivates the second definition, which introduces latent variables $Z_j^1$ and $Z_j^2$ indicating whether taxon $j$ is detected in the two samples. The second definition relies on zero-inflated models: $M_j^r | (Z_j^r = 0) = 0$, $r = 1, 2$, and it defines taxon $j$ as DA if the null hypothesis

$$H_0 : \mathbb{E}[M_j^1 | (Z_j^1 = 1)] = \mathbb{E}[M_j^2 | (Z_j^2 = 1)]$$

is rejected. The ZINB-GLM method is based on this definition. A drawback of this definition is that many observations of $M_j^1$ and $M_j^2$ would not be used for testing this hypothesis, if their corresponding $Z_j^1$ and $Z_j^2$ are inferred as zeros, resulting in a power loss. To relieve this issue, imputation can be used to rescue the likely false zeros and infer their actual values, and mbImpute achieves this by borrowing information from similar samples, similar taxon, and sample covariates, leading to the third definition of DA taxa. Assuming that the imputation is successful, we denote $M_j^{\text{imp1}}$ and $M_j^{\text{imp2}}$ as taxon $j$'s imputed counts, and $Y_j^{\text{imp1}}$ and $Y_j^{\text{imp2}}$ as the imputed abundances on the logarithmic scale, in the two samples. Then the third definition calls taxon $j$ DA if the null hypothesis

$$H_0 : \mathbb{E}[M_j^{\text{imp1}}] = \mathbb{E}[M_j^{\text{imp2}}] \text{ or } H_0 : \mathbb{E}[Y_j^{\text{imp1}}] = \mathbb{E}[Y_j^{\text{imp2}}]$$

is rejected. This third definition is advantageous in that (1) compared with the first definition, it is less affected by the existence of false zeros, whose different proportions in the two subject groups may lead to false positive DA taxa (i.e., the taxa whose non-zero counts do not exhibit a clear difference between the two groups), and (2) compared with the second definition, it uses all the observations of taxon $j$ for testing, leading to an increase in statistical power. In our implementation of imputation-empowered test, we used $H_0 : \mathbb{E}[Y_j^{\text{imp1}}] = \mathbb{E}[Y_j^{\text{imp2}}]$ because we observed that the imputed abundances exhibit bell-shaped distributions on the logarithmic scale, supporting the use of a two-sample $t$ test. If we opted for the null hypothesis $H_0 : \mathbb{E}[M_j^{\text{imp1}}] = \mathbb{E}[M_j^{\text{imp2}}]$ on the count scale, we could easily apply the Wilcoxon rank-sum test to the imputed counts.

#### Literature evidence for the selected genera in the T2D and CRC real data studies

For the analysis of T2D data, we focused on four genera, namely **Streptococcus**, **Lactobacillus**, **Clostridium**, and **Actinomyces**, which have been found related to T2D. Since obesity has long been regarded as an important risk factor for diabetes, it is reasonable to assume that enriched microbes in obesity are likely markers for diabetes as well (Nguyen *et al.*, 2011). Below is the literature evidence.

- Casarin *et al.* (2013) employed 16s rRNA sequencing on $12$ T2D and $11$ non-T2D subjects and reported a higher abundance of **Streptococcus** and **Actinomyces** in T2D subjects.
- Semova *et al.* (2012) used a zebra fish model and Ley *et al.* (2005) used a mouse model to study the relationship between gut microbiome and obesity, and they both discovered the enrichment of the Firmicutes phylum, which includes **Streptococcus**, **Lactobacillus**, and **Clostridium**, in the obesity subjects.
- Remely *et al.* (2013) applied 16s rRNA sequencing to the fecal samples of $24$ insulin-dependent T2D patients, $14$ obese participants and $19$ lean controls. They concluded that T2D patients tend to have higher proportions of lactic acid bacteria (LAB) including **Streptococcus** and **Lactobacillus**. They also reported that the **Clostridium** genus, belonging to the Firmicutes phylum, is the most abundant bacterial group detected in their experiment.

- Larsen *et al*. (2010) observed significantly higher levels of the **Lactobacillus** genus in the obesity human adults than in the control.
- Yang *et al*. (2019) performed 16s rRNA sequencing on $647$ obese and $969$ non-obese individuals to study the association between oral microbiome and obesity in African-American populations. They found that **Actinomyces** is significantly associated with increased obesity prevalence.

For the analysis of CRC data, we focused on five genera, **Fusobacterium**, **Peptostreptococcus**, **Prevotella**, **Gemella**, and **Streptococcus**, all of which have been previously reported as enriched in CRC. Below is the literature evidence.

- Allen and Jobin (2014) reviewed the literature of CRC-associated human gut microbiome, and they reported that the **Fusobacterium** genus has overabundance in tumors compared to normal tissues. They also pointed out that **Streptococcus** and **Peptostreptococcus** are enriched in luminal compartments of CRC patients in two separate Chinese cohorts (Wang *et al*., 2012; Wu *et al*., 2013).
- Dulal and Keku (2014) highlighted known microbiome risk factors for CRC and pointed out that **Fusobacterium** and **Streptococcus** are CRC-enriched.
- Nakatsu *et al*. (2015) applied 16s rRNA sequencing to $52$ CRC and $61$ non-CRC samples and reported an overabundance of **Fusobacterium**, **Peptostreptococcus**, and **Gemella** in CRC patients.
- Sobhani *et al*. (2011) applied 16s rRNA sequencing to $60$ CRC and $119$ non-CRC samples and reported **Prevotella** to be enriched in CRC. They concluded that this genus is a strong marker for CRC detection from stool samples.

#### Supplementary figures

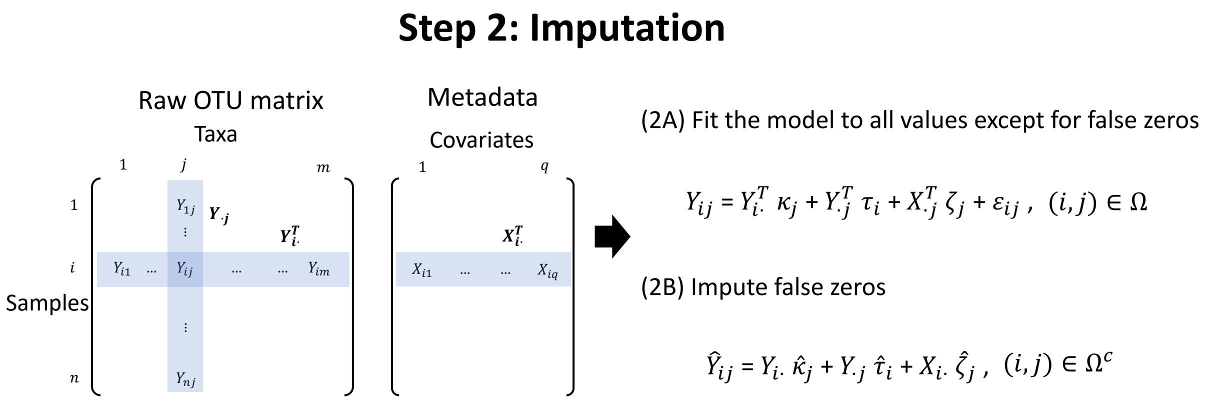


**Figure S1: A diagram illustrating the step 2 of mbImpute.** After step 1 that identifies likely false zeros, mbImpute borrows information across taxa, samples and sample covariates to impute these likely false zeros. For details, see Methods in the main text.

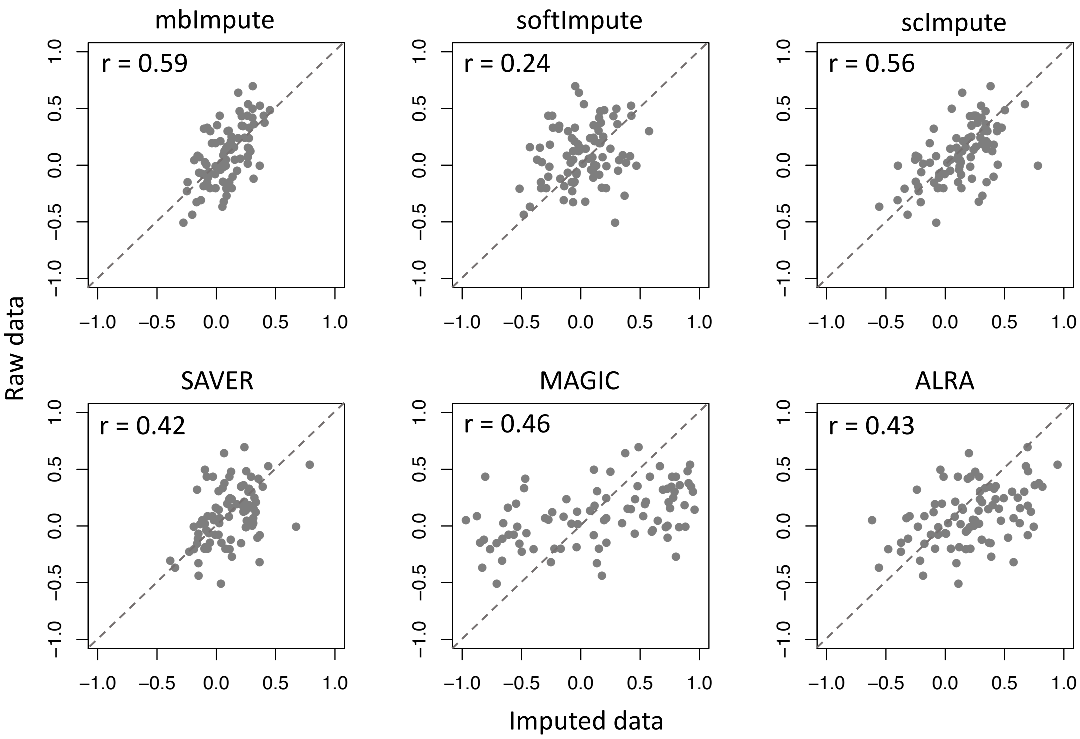


**(a)** CRC samples (Feng *et al.*, 2015)

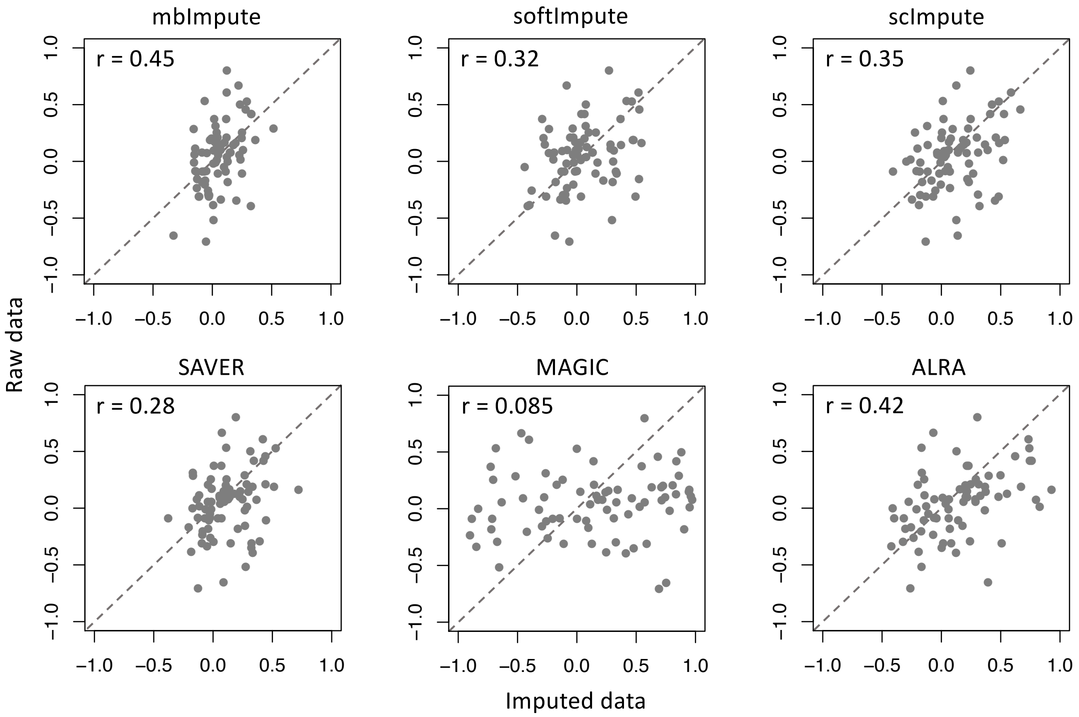


**(b)** Control samples (Feng *et al.*, 2015)

**Figure S2: Correlations between phylogenetically closely related taxa in the raw data vs. those in the imputed data.** For each pair of taxa connected by a path shorter than $4$ edges in the phylogenetic tree, we computed their correlation in the raw data using their mutual non-zero abundances (vertical axis) and their correlation in the imputed data using all the abundances (horizontal axis). The Pearson correlation between these two sets of correlations is reported for each imputed dataset, showing that mbImpute achieves the highest correlation in both (a) the CRC samples and (b) the control samples.

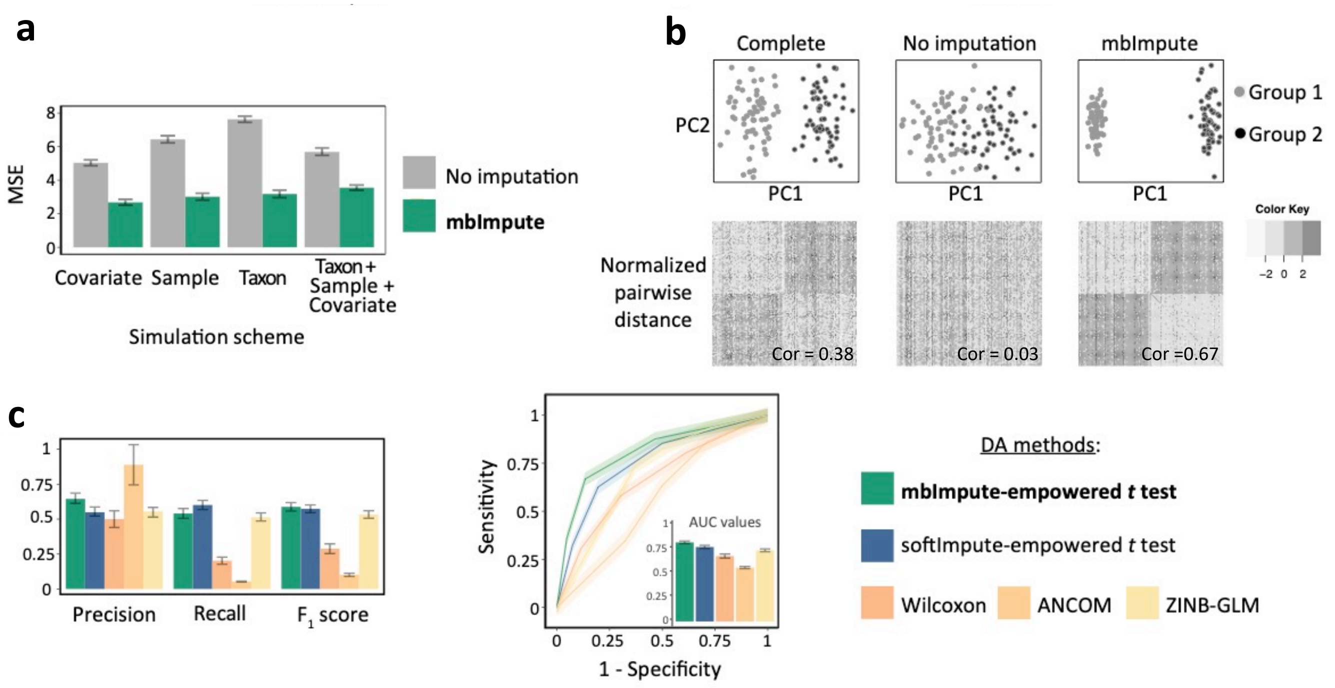


**Figure S3: mbImpute enhances DA taxa detection in simulated WGS datasets.(a)** MSE between the complete data and the zero-inflated data or the imputed data by mbImpute under four simulation schemes in Simulation 3. **(b)** Data in Simulation 4. Top: the complete data, the zero-inflated data, and the imputed data by mbImpute in the first two principal components (PC1 and PC2). Bottom: normalized pairwise Euclidean distances between samples are shown for each dataset; each row of the distance matrix was normalized to have zero mean and unit standard deviation. The correlations between the perfect sample separation and distance matrices is labeled on the right corner of each heatmap. **(c)** DA taxon detection results in Simulation 4. Left: precision, recall and $F_1$ scores of three existing DA methods (Wilcoxon rank-sum test, ANCOM, and ZINB-GLM), and two imputation-empowered $t$ test (softImpute and mbImpute). Right: ROC curves and corresponding AUC values of the five methods.

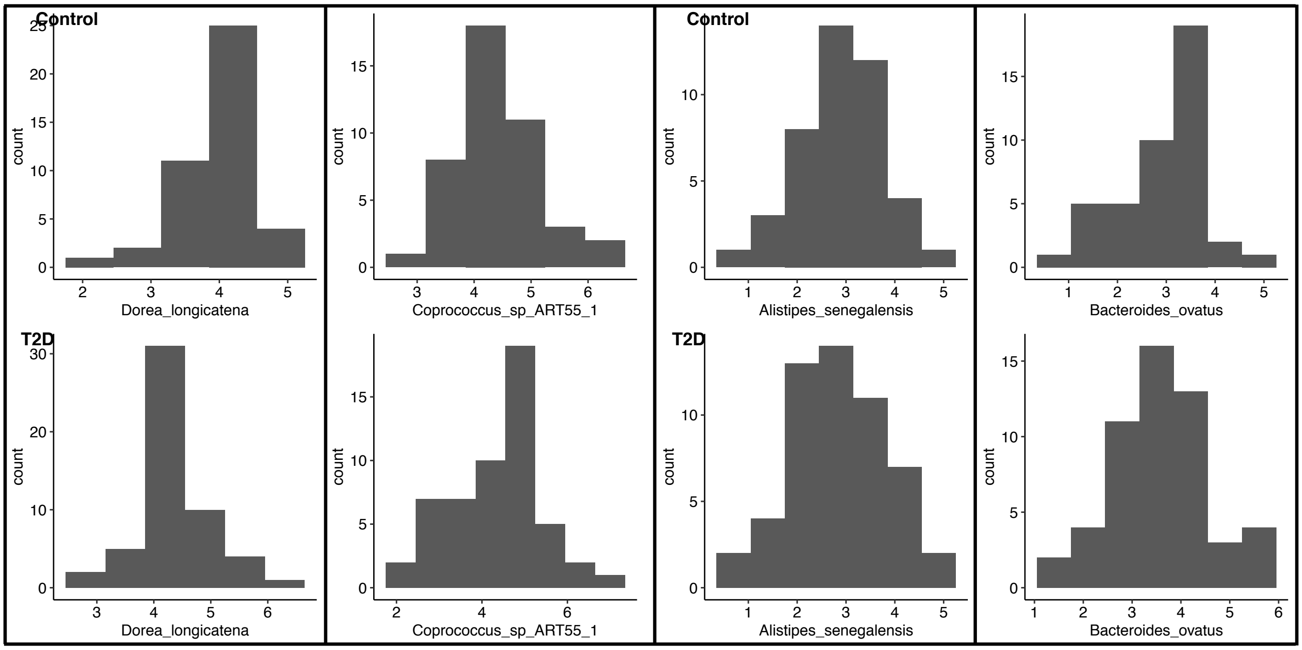


**Figure S4: Distributions of four randomly chosen taxa from Karlsson *et al.* after mbImpute is applied.** The imputed abundances are approximately normal.

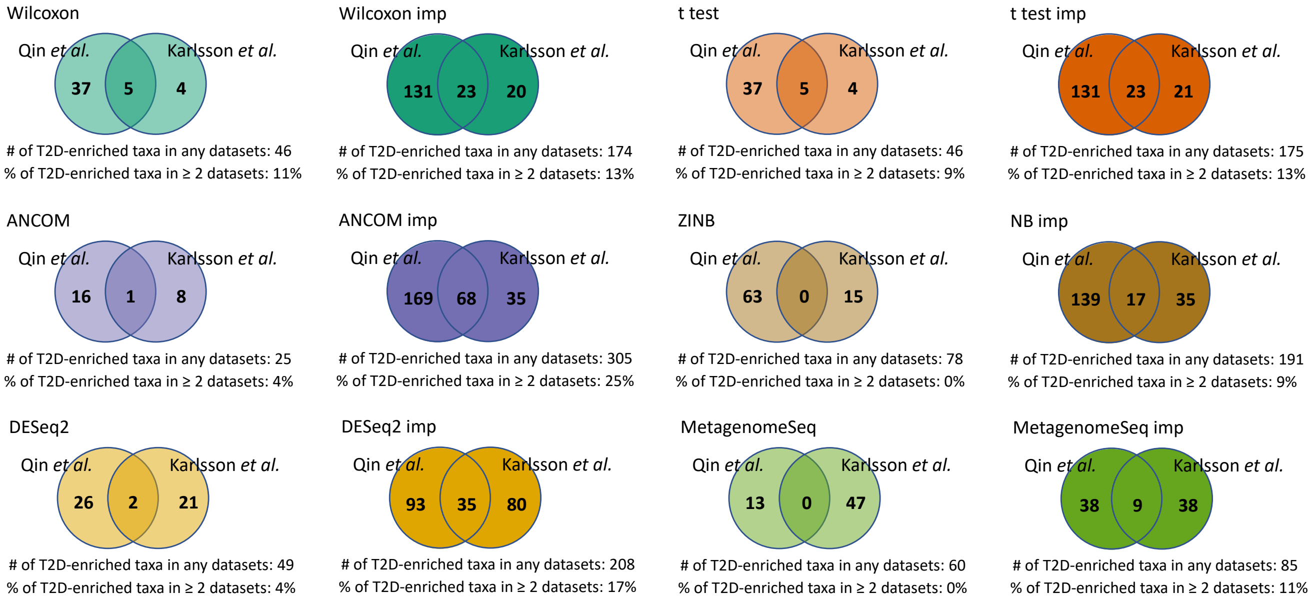


**Figure S5: mbImpute increases the power and reproducibility of DA taxon identification in two T2D-related real datasets from Qin *et al.*, and Karlsson *et al.*** Venn diagrams showing the overlap of T2D-enriched taxa in two datasets identified by each of six DA methods (Wilcoxon rank-sum test, $t$-test, ANCOM, ZINB/NB-GLM, DESeq2, and metagenomeSeq) before (lighter color) and after mbImpute (darker color). The number of DA taxa identified in either dataset and the percentage of DA taxa identified in both datasets—100(# DA taxa identified in both datasets / # DA taxa identified in either dataset)%—are shown below each Venn plot.

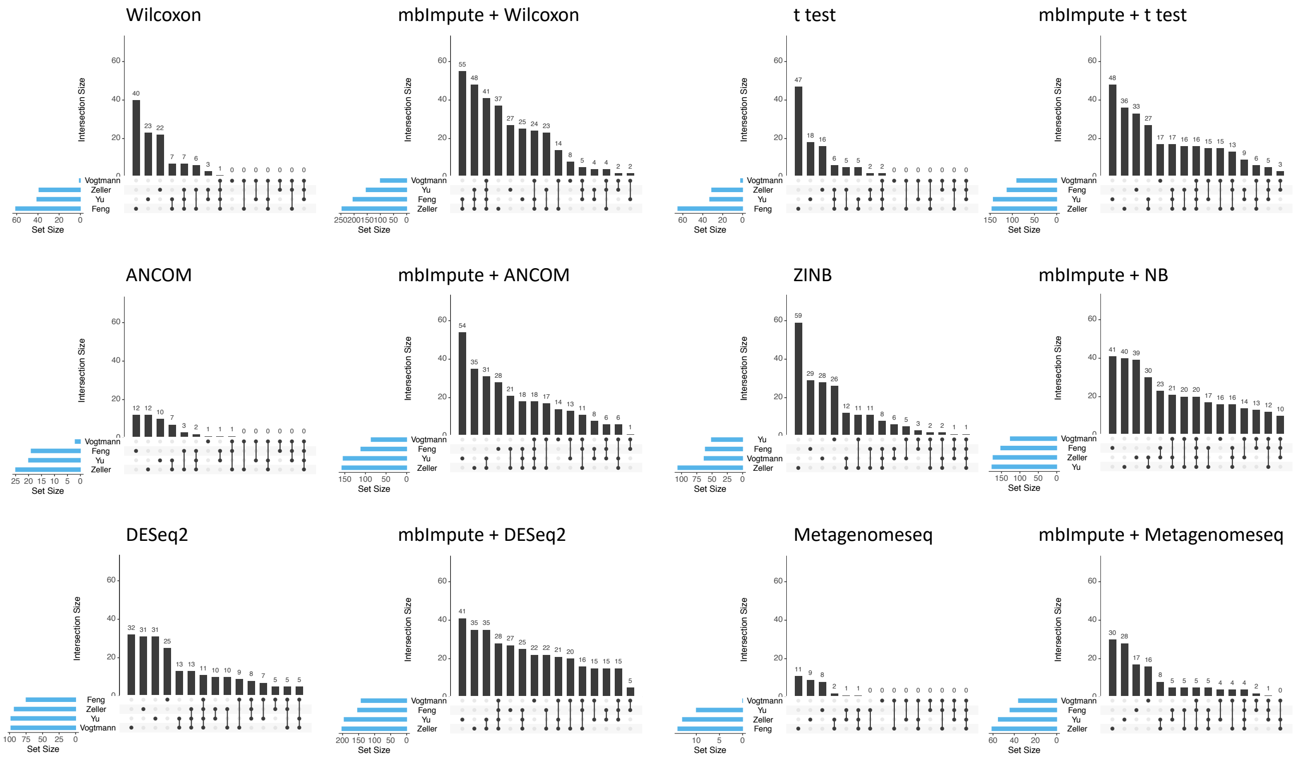


**Figure S6: mbImpute increases the power and reproducibility of DA taxon identification in four CRC-related real datasets from Zeller *et al.*, Feng *et al.*, Vogtmann *et al.*, and Yu *et al.*** Diagrams showing the overlap of CRC-enriched taxa in four datasets identified by each of six DA methods (Wilcoxon rank-sum test, $t$-test, ANCOM, ZINB/NB-GLM, DESeq2, and metagenomeSeq) before and after mbImpute.

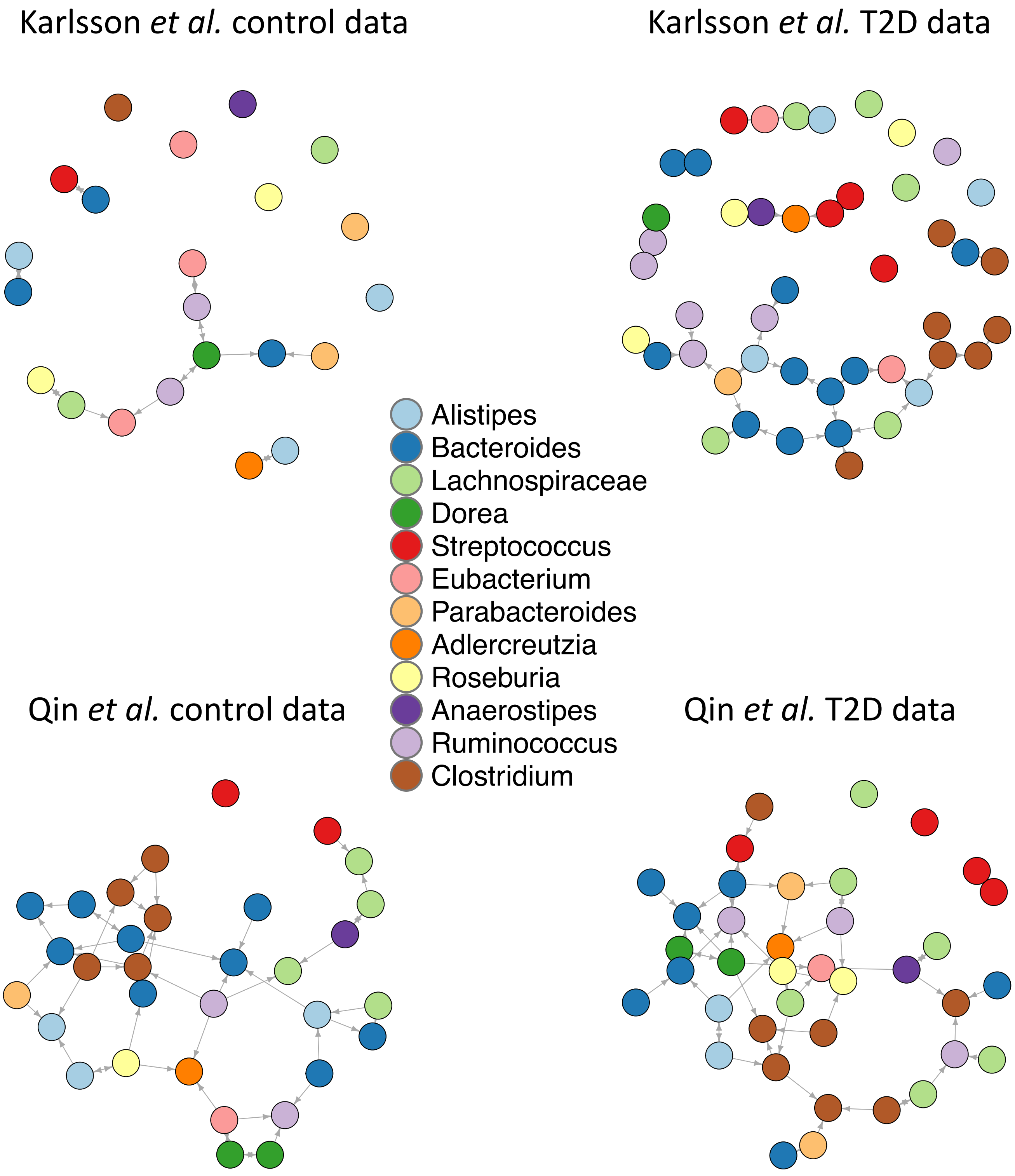


**Figure S7: Visualization of genus-level taxon interaction networks constructed from two T2D datasets after mbImpute.** Four genus-level interaction networks constructed by the PC algorithm (Kalisch *et al.*, 2007) on the Karlsson *et al.* and Qin *et al.*'s control samples and T2D samples after mbImpute.

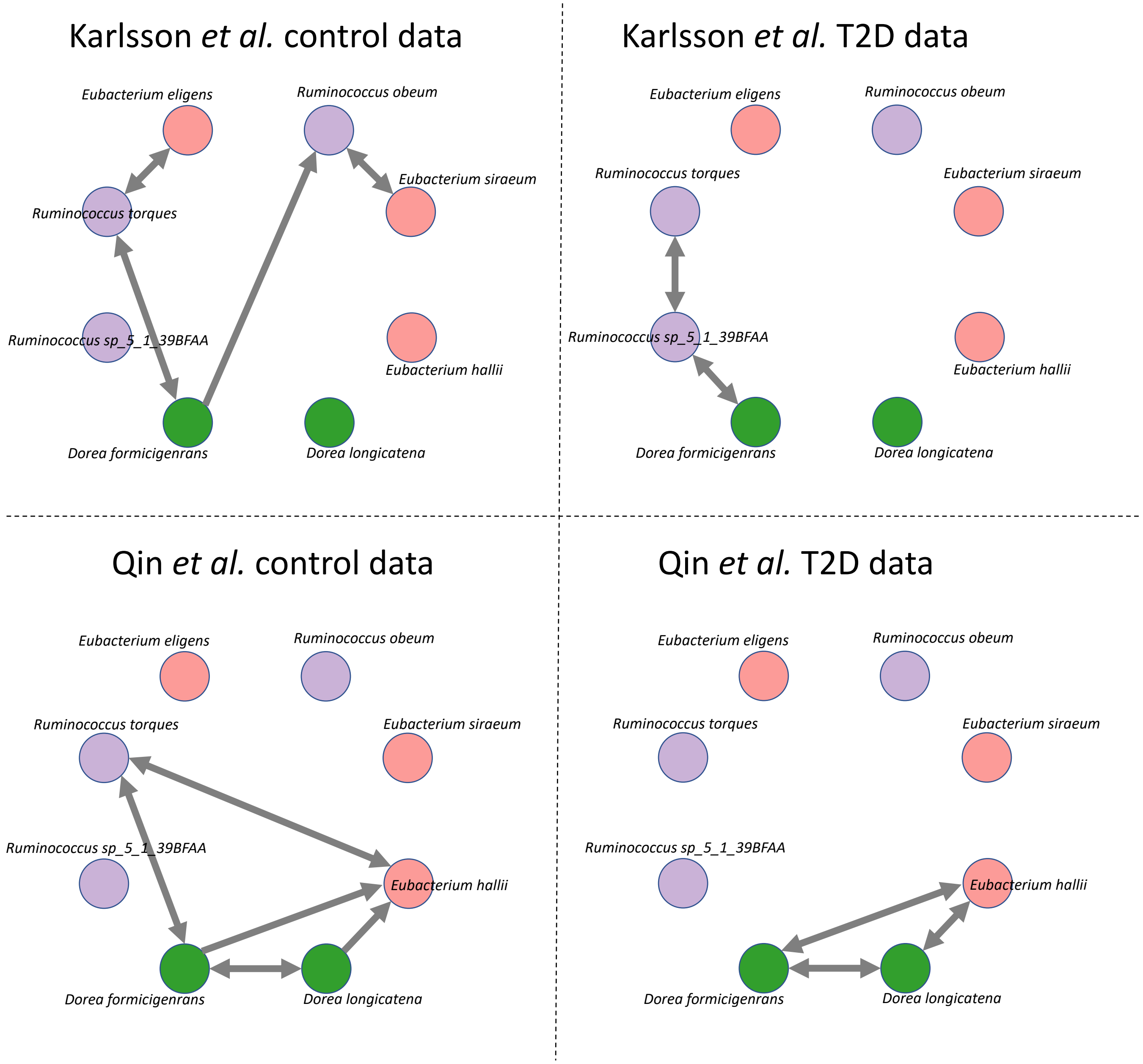


**Figure S8: Visualization of species-level taxon interaction networks of three genera (*Eubacterium*, *Ruminococcus*, and *Dorea*) from two T2D datasets after mbImpute.** Four strain-level interaction networks constructed by the PC algorithm on the Karlsson *et al.* and Qin *et al.*'s control samples and T2D samples after mbImpute.

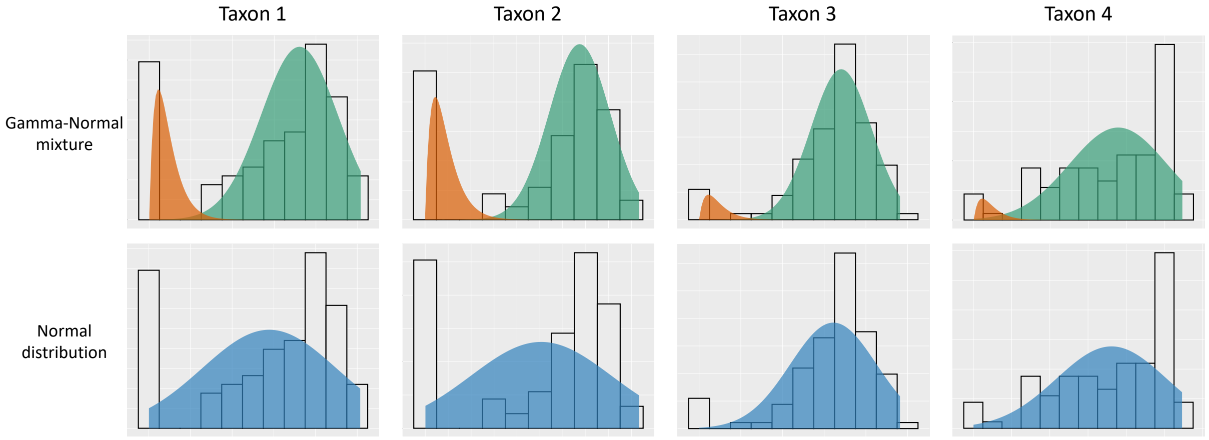


**Figure S9: Abundance distributions of four taxa in the data from (Zeller *et al.*, 2014).** The same histogram is displayed in each column to show each taxon's abundance distribution. Top panel: fitted Gamma-Normal mixture model, where the red and green areas represent the Gamma and Normal components, respectively. Bottom panel: fitted Normal distribution represented by the blue area. For details, see Methods in the main text.

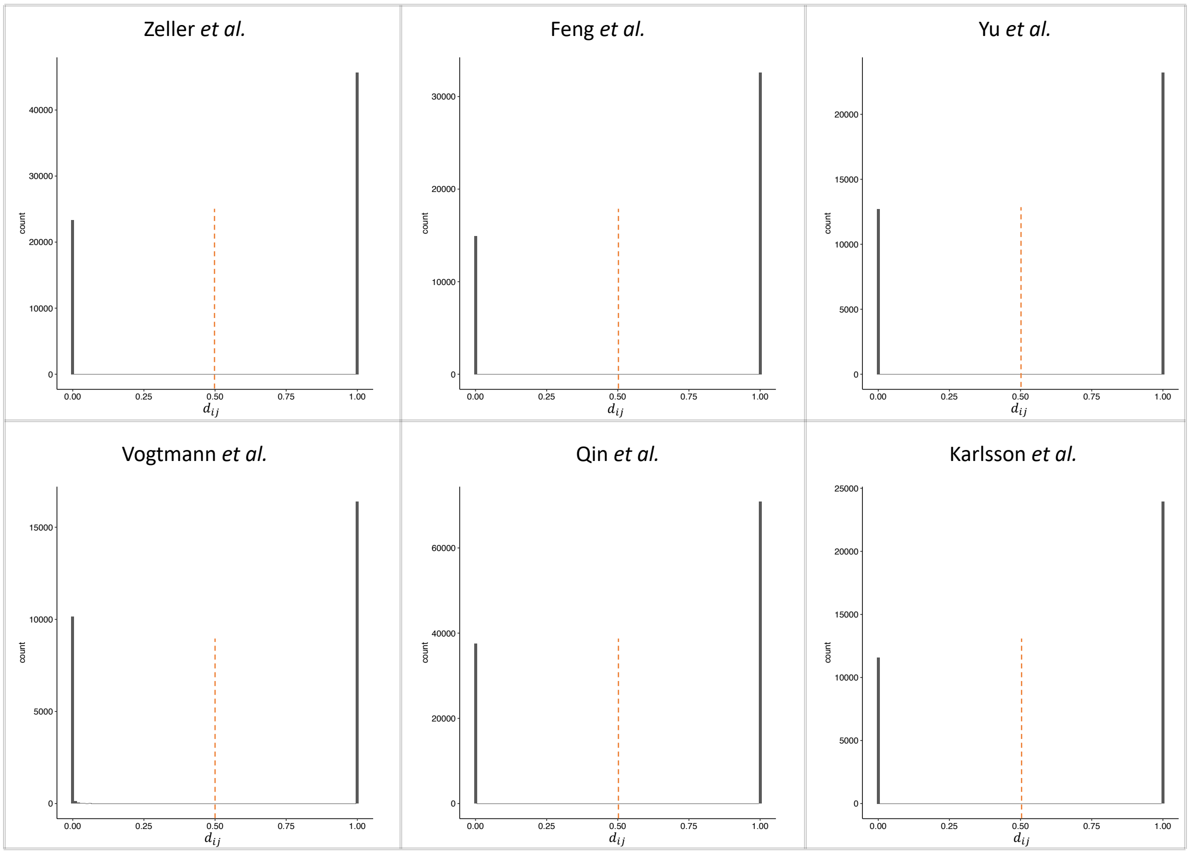


**Figure S10: The distributions of $d_{ij}$'s calculated in step 1 of mbImpute applied to six real datasets.** The 0.5 threshold is indicated by the red dotted line. For details, see Methods in the main text.
